# DeDoc2 identifies and characterizes the hierarchy and dynamics of chromatin TAD-like domains in the single cells

**DOI:** 10.1101/2022.08.23.505046

**Authors:** Angsheng Li, Guangjie Zeng, Haoyu Wang, Xiao Li, Zhihua Zhang

## Abstract

Topologically associating domains (TAD) are functional chromatin units with hierarchical structure. However, the existence, prevalence and dynamics of such hierarchy in single cells remain unexplored. Here, we report a new generation TAD-like domain (TLD) detection algorithm, named deDoc2, to decode the hierarchy of TLDs in single cells. With dynamic programming, deDoc2 seeks genome partitions with global minimal structure entropy for both whole and local contact matrix. Compared to state-of-the-art tools, deDoc2 can uniquely identify the hierarchy of TLDs in single cells, in addition to outperforming its competitors. By applying deDoc2, we showed that the hierarchy of TLDs in single cells is highly dynamic during cell cycle, as well as among human brain cortex cells, and that it is associated with cellular identity and functions. Thus, our results demonstrated the abundance of information potentially encoded by TLD hierarchy for functional regulation. The deDoc2 can be freely accessed at https://github.com/zengguangjie/deDoc2.

## Background

The eukaryotic genome has a hierarchical configuration ^1,2^, as revealed by imaging technologies^3^ and chromosome conformation capture (3C)-based technologies ^4-12^, e.g., Hi-C ^8^. These configurations, including chromosomal territories ^8,13,14^, A and B compartments ^8^, topologically associating domains (TADs) ^14,15^, compartment domains ^16^, or CTCF loop domains ^17^, and chromatin loops ^17-19^, have been routinely discussed ^2^. TADs may be one of the most investigated chromatin features in the literature. Studies have found that TADs are also organized in hierarchical fashion, e.g., the hierarchy of domains-within-domains (metaTAD) through TAD-TAD interactions at the large scale ^20,21^. Hierarchical organization of TADs is functionally significant in that the hierarchical levels of metaTAD correlate with key epigenomic and expression features, and smaller sub-TADs are specifically associated with gene regulation ^22-24^. A long list of hierarchical TAD detection tools is available in the literature, such as Armatus ^25^, TADtree ^26^,GMAP ^27^,IC-Finder ^28^,PSYCHIC ^29^,CaTCH ^30^,3DNetMod ^31^,Matryoshka ^32^,deDoc ^33^,OnTAD ^34^,SpectralTAD ^35^,TADpole ^36^,HiCKey ^37^, and SuperTAD ^38^. However, all these tools were based on bulk Hi-C data.

TAD-like domains (TLDs) have been found in single cells^39^. Tools for single-cell TLD detection have been rapidly developed since the emergence of single-cell Hi-C technologies ^40^, including scHiCluster ^41^,deTOKI ^42^, and Higashi ^43^. Some tools involved in data imputation were needed to solve the sparseness of single-cell Hi-C, e.g., scHiCluster ^41^ or Higashi ^43^, while deTOKI^42^ utilized non-negative matrix factorization (NMF) to address the problem. The advantage of NMF lies in its low rank representation, which retrieves key information embedded in the noisy sparse data. As a sparse non-negative matrix, the sparsity of scHi-C data can also be solved by NMF. However, all existing methods mainly utilized local information or arbitrarily borrowed information from neighboring cells, making it hard to explore higher-level structures. Nevertheless, the existence of TAD-like domains signaled the likelihood of higher-level chromatin configuration in single cells not detectable by the above tools.

Given the large cell-to-cell variations of chromatin architecture observed in individual cells, the existence of hierarchical TLDs remains a non-trivial question. In other words, the hierarchy of TADs observed in bulk Hi-C could be a completely, or partially, emergent property of the cell population. That is, the dynamics of chromatin in single cells *per se* may generate, at least in part, the hierarchy ^20,21^. Since the origin and dynamics of TADs are keys to understanding gene regulation ^21,44^, it is essential to unravel the hierarchical nature of single-cell TLD. However, a systematic survey of TLD hierarchy, including its dynamics, in single cells remains a major challenge in the field.

We recently developed deDoc ^33^, a tool which detects the hierarchical TAD structure with sparse Hi-C data. The deDoc was based on the structural information theory ^45^, which measures the uncertainty embedded in the dynamics of a graph. Minimizing structural entropy is an intuitive way to decode the essential structure of a graph in which perturbations caused by random variation and noise have been reduced to a minimum. Thus, deDoc integrates the whole information in the graph, making the algorithm robust against data noise and sparseness. Recently, a tool named SuperTAD, which is based on structural entropy, also appeared in the literature. It is dedicated to the identification of hierarchical TADs using global minimization of structural entropy ^38^. However, neither deDoc nor SuperTAD work properly with single-cell Hi-C data.

Here, we present a new generation of TLD detector, deDoc2, which can reliably predict hierarchical TLD structures in single cells by dynamic programming. Compared to state-of-the-art tools, deDoc2 not only outperforms its competitors in reliable TLD identification in single cells, but it is also robust to various data imputation algorithms. Using deDoc2, we found that hierarchical TLDs prevalent in single cells were dynamic, experiencing changes during cell cycle. We also found that TLD hierarchy was remarkably different among human brain prefrontal cortex cells. Here, we present examples, showing that both structure and hierarchy of TLDs may be subject to functional regulation in single cells. In short, deDoc2 opens the door to a systematic understanding of TLD hierarchy and, as such, it provides a new dimension by which to decipher the dynamics of single-cell 3D chromatin structure.

## Results

### The deDoc2 is a single-cell hierarchical TAD-like domain (TLD) predictor

We developed a deDoc2 package to detect hierarchical TLDs with single-cell Hi-C data (Fig. 1). Similar to our previously developed deDoc ^33^, deDoc2 seeks to partition the genome into domains with minimal structural entropy. Briefly, the Hi-C contact map was regarded as a network-connected matrix, and the global optimal partition of the network with minimal two-dimensional structural entropy was detected by a dynamic programming algorithm (Fig. 1). However, deDoc2 goes beyond deDoc with its dynamic programming algorithm used to optimize two-dimensional, but not high-dimensional ^38^, structural entropy to achieve global optimization (Fig. 1a). This allows a minimalist approach without losing key information.

**Figure 1.**
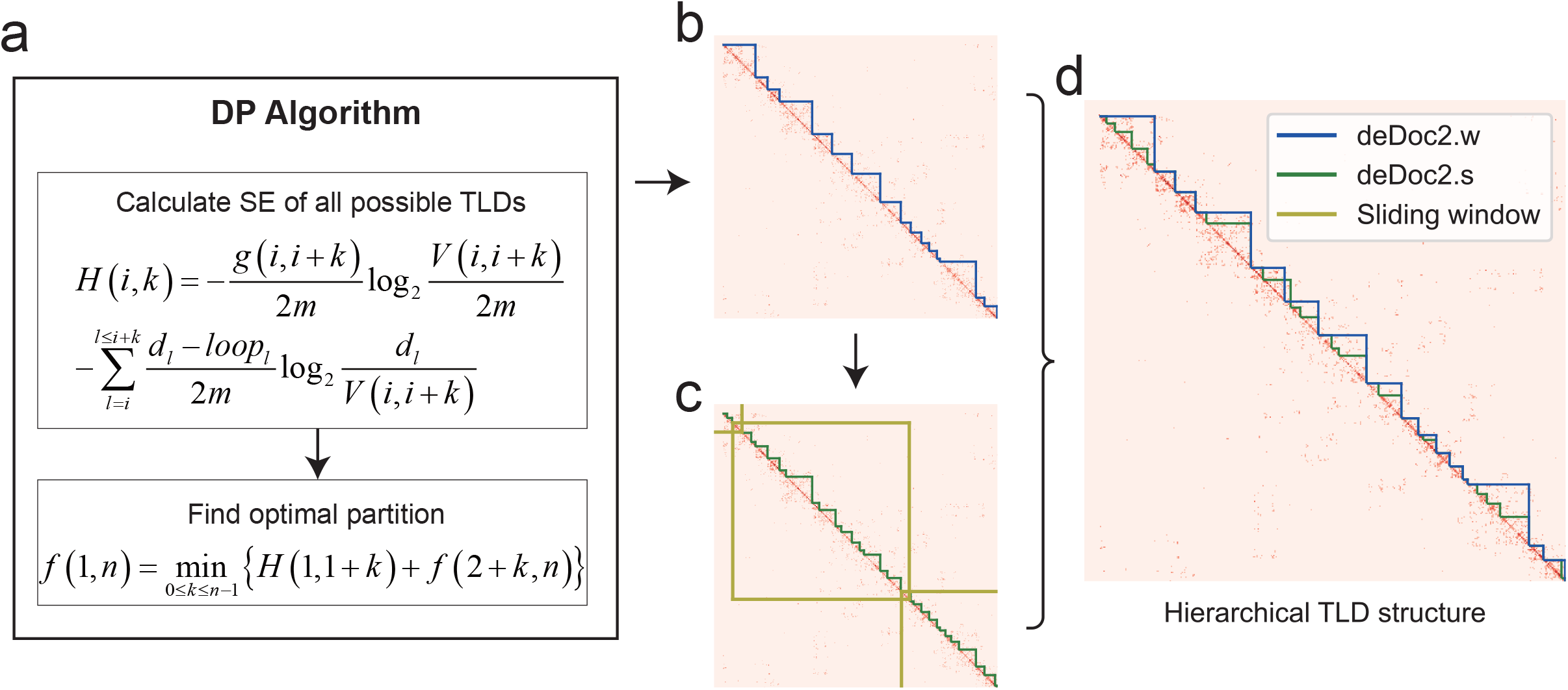
Flow chart of deDoc2. **a**. Dynamic programming algorithm used to minimize two-dimensional structural entropy, which includes calculating structural entropy of all possible TLDs and finding optimal partition with recurrence relation. **b**. Example of TLDs called by deDoc2.w, Hi-C data from Rao et al., cell 14 chromosome 18, 18Mb∼34Mb. We minimize two-dimensional structural entropy of whole Hi-C to obtain the TLDs. **c**. Example of TLDs called by deDoc2.s, Hi-C data from Rao et al., cell 14 chromosome 18, 18Mb∼34Mb. We minimize two-dimensional structural entropy of Hi-C in the sliding window to obtain TLDs. **d**. Hierarchical TLD structure consisting of TLDs called by deDoc2.w and deDoc2.s.

The deDoc2 package consists of two predictors, deDoc2.w and deDoc2.s, to predict higher and lower level TLDs (Fig. 1b, c), respectively. The deDoc2.w minimizes structural entropy of the Hi-C contact map of whole chromosomes (Fig. 1b), while deDoc2.s minimizes structural entropy in the matrices of sliding windows (10Mb) along the genome (Fig. 1c). Combining the results from the predictors, we can obtain the final two-layer nested TLDs (Fig. 1d). In addition to TLD prediction, the deDoc2 package provides a tool to determine proper binsize for the given Hi-C data. Unlike deDoc which implements this feature with one-dimensional structural entropy, deDoc2 uses normalized decoding information to optimize binsize (Fig. S1, Supplementary Information (SI)).

Next, we sought to assess the performance of deDoc2 by comparing it with seven state-of-the-art algorithms, i.e., Insulation score (IS) ^46^, deDoc^33^, SpectralTAD ^35^, GRiNCH ^47^, deTOKI ^42^, scHiCluster ^48^ and Higashi ^49^. These seven methods can be grouped into those mostly designed for sparseness (SpectralTAD and GRiNCH) or single cell (deTOKI, scHiCluster and Higashi). We first assessed them with downsampled ultra-sparse Hi-C data, followed by simulated and experimental single-cell Hi-C data. Sparsity was defined as the proportion of non-zero entries in the Hi-C matrix. The assessment was made with and without data imputation. We mainly described the results from imputation-free data, while similar performance with data imputation can be found in SI (Figs. S2 and S3).

### deDoc2 worked well with downsampled bulk Hi-C at the single-cell level

To assess the tools, we generated a series of downsampled data from ^50^ with sampling rates of 0.1%, 0.05% and 0.025% and set default binsize=40kb. The dataset at the lowest sampling rate contains about 0.23M contacts, mimicking the sequencing depths of scHi-C. For example, the median number of contacts in the data generated by Flyamer and colleagues ^51^ was 0.339M.

The deDoc2 outperformed the other tools in the following two respects. First, deDoc2 predicted TLDs more accurately than all other predictors. We took the TADs identified by full data with the predictors themselves as the gold standard and quantified the accuracy of predictions by the similarity, i.e., adjusted mutual information (AMI)^52^ and weighted similarity (WS)^33^. The predictions of deDoc2.w have the highest AMI and WS of all sample rates among all other algorithms, followed by Higashi, IS, deDoc2.s and deTOKI (Fig. 2a and b). The order of the latter 4 algorithms was not consistent between the two accuracy indices; however, all are ranked in the top. Second, deDoc2 is more robust to binsize than the other algorithms. We compared the TLDs detected based on different binsizes, i.e., 30kb *vs*. 60kb and 40kb *vs*. 80kb. Again, we found that deDoc2.w had the highest AMI and WS among all algorithms for both binsize pairs, followed by Higashi, deDoc2.s and IS. The latter 3 are also ranked in the top (Fig. 2c and d). Last, the characteristic binding of structural protein CTCF, or histone marks, was found to be more enriched in deDoc2-predicted TLD boundaries than other tools. We compared the enrichment of ChIP-seq peaks of H3K4me3, H3K36me3, and CTCF at the predicted TLD boundary regions (Fig. 2e). The enrichment of all three marks is significantly higher in the boundaries predicted by deDoc2.w than in those predicted by the other tools at all sampling rates (Fig. 2e). At the lowest sampling rate, Higashi and deDoc2.s, together with deTOKI, following deDoc2.w, have considerably high enrichment for those epigenetic marks. Taken together, our assessments suggest that deDoc2 can stably and accurately predict TLDs with ultra-low resolution Hi-C data.

**Figure 2.**
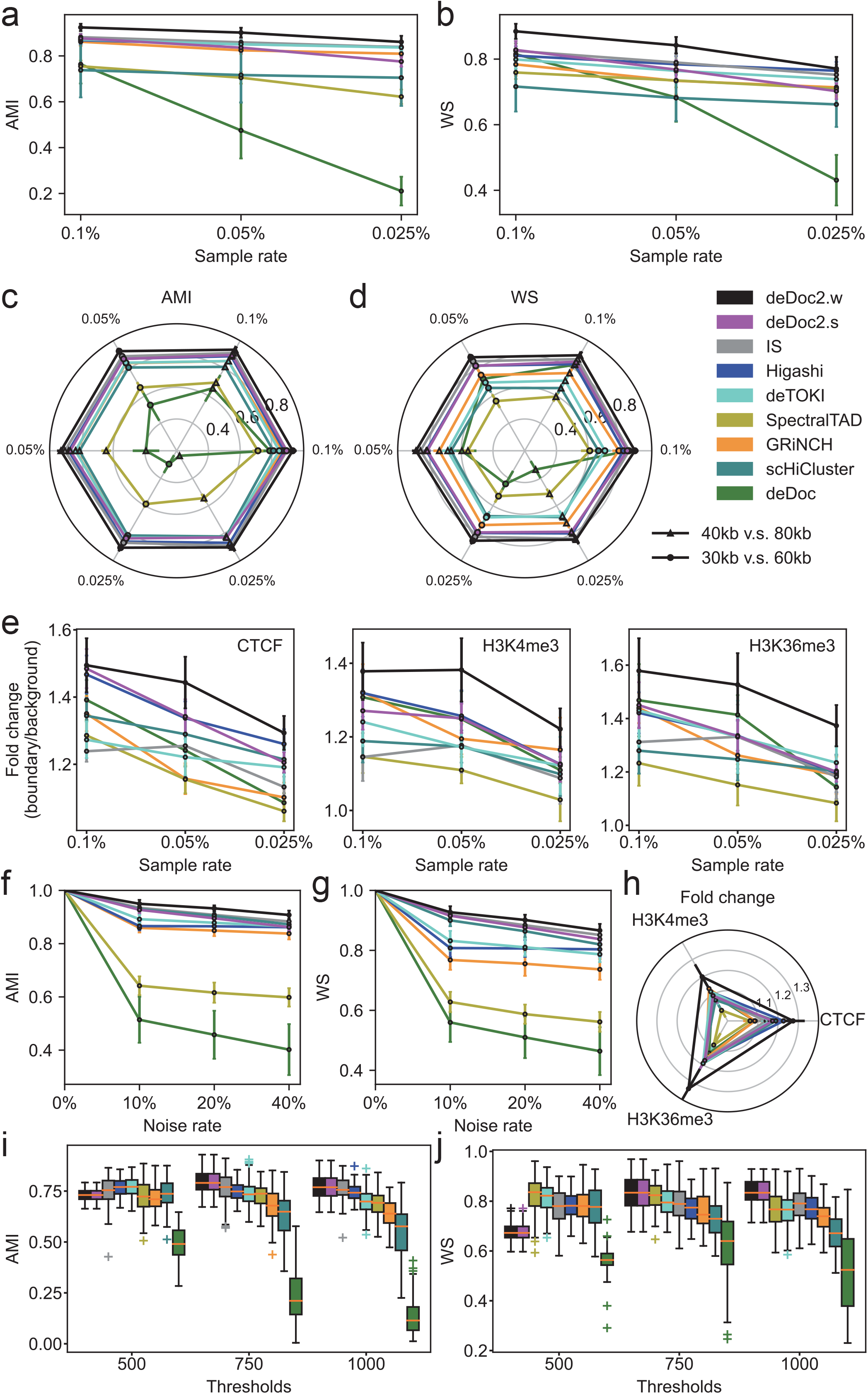
Assessment of deDoc2 with downsampled data, noisy data and simulated data based on Rao’s data. The downsampled Hi-C data come from IMR90 cell type from Rao et al. with downsample rate of 0.1%, 0.05% and 0.025%. Noisy data were generated by adding contacts at 0.025% downsampled Hi-C data with noise rate 10%, 20% and 40%. Simulated data come from chr18:50-55 Mb for GM12878 ensemble Hi-C from Rao’s data. **a**. AMI between TLDs from downsampled Hi-C at three downsample rates and bulk Hi-C called by predictors. **b**. WS between TLDs from downsampled Hi-C at three downsample rates and bulk Hi-C called by predictors. **c**. Robustness to binsize. AMI between TLDs from downsampled Hi-C of different binsize (30kb *vs*. 60kb and 40kb *vs*. 80kb). **d**. Robustness to binsize. WS between TLDs from downsampled Hi-C of different binsize (30kb *vs*. 60kb and 40kb *vs*. 80kb). **e**. Fold change of ChIP-seq peak signals (CTCF, H3K4me3 and H3K36me3) between TLD boundaries and background (regions away from boundaries). TLDs are called from downsampled Hi-C at three downsample rates. Error bars represent 95% confidence interval. **f**. AMI between TLDs from original downsampled Hi-C and noisy Hi-C at three noise rates. **g**. WS between TLDs from original downsampled Hi-C and noisy Hi-C at three noise rates. **h**. Fold change of ChIP-seq peak signals (CTCF, H3K4me3 and H3K36me3) between TLD boundaries and background (regions away from boundaries). TLDs are called from noisy Hi-C at three noise rates. Error bars represent 95% confidence interval. **i**. AMI between TLDs from simulated scHi-C at different simulation thresholds and bulk Hi-C. **j**. WS between TLDs from simulated scHi-C at different simulation thresholds and bulk Hi-C.

### deDoc2 is robust against data noise in downsampled Hi-C data

Given the ultra-sparse nature of scHi-C data, TLD predictors may be sensitive to data noise. To assess the robustness of deDoc2 against data noise, we randomly modified 10%, 20%, and 40% of entries in the Hi-C matrices with a replacement to mimic data noise in the downsampled data at the lowest sampling rate, 0.025% (see Methods). We found that deDoc2 is robust to noise because both AMI and WS of TLDs between noisy and original downsampled data decreased only slightly when noise rate increased (Fig. 2f, 2g). Among all TLD predictors, deDoc2.w achieved the highest AMI and WS, followed by IS, deDoc2.s and scHiCluster. The deDoc2.w achieved enrichment much higher than all other predictors, and Higashi, scHiCluster and deDoc2.s all achieved quite high enrichment at all noise rates (Fig. 2h).

### deDoc2 worked well with simulated scHi-C data

To assess the effect of cell-to-cell variance in the cell population to the performance of TLD predictors, we tested the predictors with simulated scHi-C data. The simulation process was conducted on the basis of our previous work^42^. To simplify the simulation, the structures in the ensemble were assumed to be evenly distributed in the cell population, and a 5Mb-long genome region, i.e., chr18:50-55Mb, was randomly chosen as an example. As 5Mb is smaller than the default sliding window size for deDoc2.s, the results for deDoc2.s and deDoc2.w were, in fact, identical, but we only reference deDoc2 in this section. Threshold (*D*) occurs in the simulation process (see Methods). We tested *D*s with 500, 750 and 1000, representing 20%, 40% and 60% quantiles, respectively (Fig. 2i, 2j). We simulated Hi-C data from a 3D chromosome structure ensemble containing about 100 single cells ^42^

The deDoc2 could accurately predict domain structures in simulated scHi-C data. We generated simulated scHi-C data about 1000 and 0.35M Hi-C contacts for each single cells and the reference Hi-C in that 5M-long chromosome region, respectively. We compared accuracies (AMI and WS) of the predictions to the reference. For D=750 and 1000, deDoc2 predicted the most accurate TLDs compared to the reference based on the highest AMI and WS (Fig. 2i and j). For D=500, deDoc2, together with other tools, except deDoc, predicted TLDs with almost indistinguishable AMI; however, the prediction of deDoc2 resulted in a substantially smaller WS compared to the other tools. This may have been caused by the bias to local contact of Hi-C matrix when D=500. In this case, deDoc2 is prone to separate the large domains into smaller ones, leading to a larger number of predicted TLDs compared to the other tools (Fig. S4a). However, this issue can be resolved by data imputation (Fig. S3b-c, S4b), which is a common practice in single-cell data analysis^21^. In general, the performance of deDoc2 can be slightly improved by proper data imputation (see SI). Taken together, deDoc2 performed well in most cases in the simulated scHi-C data.

### deDoc predicts TLDs well with experimental scHi-C data

Next, we compared predictions among deDoc2, deDoc^33^, SpectralTAD^35^, GRiNCH^47^, deTOKI^42^, scHiCluster^48^ and Higashi^49^, using three representative experimental scHi-C datasets (hereinafter denoted as Tan’s ^53^, Nagano’s ^54^, and Lee’s ^55^) (Supplementary Table 1-3). We found the predictions of deDoc2, in most cases, to be among the top category.

As absent a gold standard for TLDs in single cells, we assessed performance indirectly. First, deDoc2 predicted TLDs with lower structural entropy and higher modularity (Fig. 3a and b). Structural entropy and modularity are two network properties that have been used to infer TADs for bulk Hi-C data ^31,33^, i.e., a better defined TAD will have smaller structural entropy^33^ and larger modularity^31^. We tested the structural entropy and modularity of TLDs on 16 GM12878 cells from Tan’s dataset and 20 randomly picked mouse ES cells from Nagano’s dataset. Predictions of deDoc2.w and deDoc2.s had the first and second smallest structural entropies, respectively, in both datasets (Fig. 3a, Fig. S4c). For modularity, the prediction of deDoc2.w had the largest modularity in both datasets, while deDoc2.s-predicted TLDs were similar to those of the other tools (Fig. 3b, Fig. S4d). Second, CTCF and histone marks were enriched at the boundary regions of TLDs predicted by deDoc2, Higashi and GRiNCH in experimental scHi-C data (Fig. 3c, Fig. S4e). For all marks in both datasets, none of the tools showed enrichment of predicted TLD boundaries. The prediction of deDoc2.w showed the highest enrichment among all three marks in at least one dataset. Higashi’s prediction showed the highest enrichment for CTCF and H3K4me3 in Nagano’s. GRiNCH’s prediction showed the highest enrichment for H3K4me3 and H3K36me3 in Tan’s. Even in cases for which the predictions of deDoc2 did not show the highest enrichment, it could also be ranked at top category by having enrichment comparable to highest tool (Fig. 3c, Fig. S4e).

**Figure 3.**
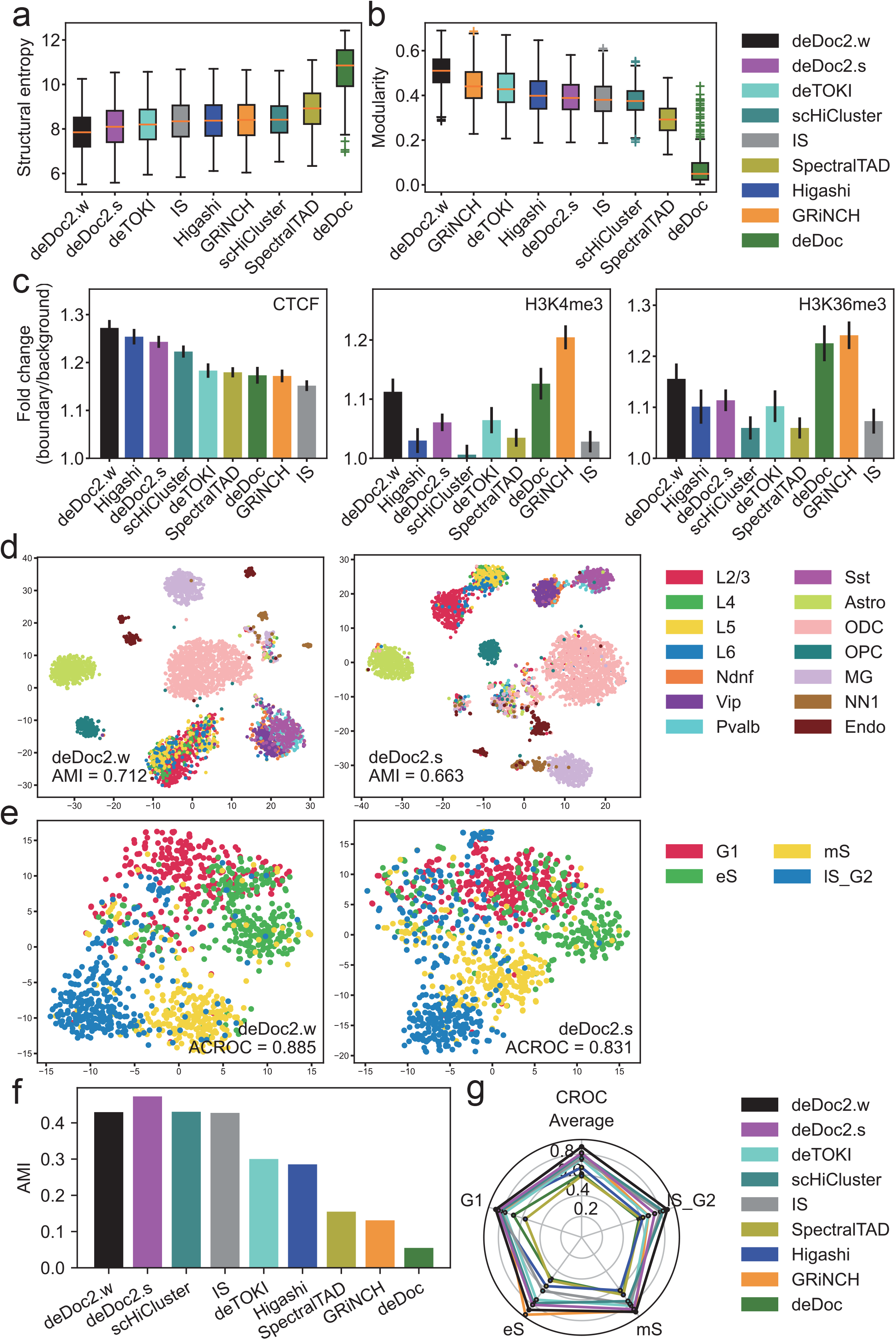
Assessment with experimental scHi-C data. We assessed deDoc2 with datasets from Tan et al., Nagano et al. and Lee et al. **a**. Structural entropy of predicted TLDs in Tan’s data. **b**. Modularity of predicted TLDs in Tan’s data. **c**. Fold change of ChIP-seq peak signals (CTCF, H3K4me3 and H3K36me3) between TLD boundaries and background (regions away from boundaries). TLDs are called from Tan’s data. Error bars represent 95% confidence interval. **d**. Cell embedding of TLDs predicted by deDoc2 with RWR imputation from humanPFC cells in Lee’s data (4238 cells). **e**. Cell embedding of TLDs predicted by deDoc2 with RWR imputation from mES cells in Nagano’s data (1137 cells). **f**. AMIs of embedding of TLDs from humanPFC cells in Lee’s data (560 cells). **g**. CROCs of embedding of TLDs from mES cells in Nagano’s data (400 cells).

Third, embedding scHi-C data with deDoc2-predicted TLDs results in satisfactory cell clustering (Fig. 3d-g). We applied the eight algorithms on Nagano’s and Lee’s datasets and took TLD boundaries as text input. Then, we performed dimensionality reduction on their term-frequency-inverse document frequency (TF-IDF) values using TrunctedSVD ^56^. The last embedding could be visualized by t-SNE (Fig. 3d-g). In Lee’s dataset, 4238 single cells are from the human prefrontal cortex (PFC) and previously identified as having 14 cell types with CpG methylation levels^55^. Consistent with clustering in the original work, the non-neuronal brain cell types, i.e., Astro, ODC, OPC and MG, can be clearly distinguished by both deDoc2.w and deDoc2.s (Fig. 3d). In addition, clustering revealed two brain neuron subtype-clusters, i.e., excitatory neuron subtypes, including L2/3, L4, L5 and L6, and inhibitory neuron subtypes, including Pvalb, Sst, Ndnf and Vip. These two suptype-clusters could not be clustered by chromatin contact alone in the original work^55^. We used AMI to quantify the embedding. The deDoc2.s had the highest AMI value on Lee’s data, followed by scHiCluster, deDoc2.w and IS (Fig. 3f). In Nagano’s dataset, single-cell Hi-C data come from four cell cycle stages^54^, i.e., “G1” phase, “early-S” phase, “mid-S” phase, “late-S/G2” phase, and “2n DNA” stages. As we failed to identify a sufficient number of TLDs from “2n DNA” stage, we excluded those data from subsequent analysis. After quality control, a total of 1171 cells were analyzed. The visualization showed a clear circular cell embedding with G1, eS, mS and lS_G2 cells clustering in a clockwise manner for both deDoc2.w and deDoc2.s (Fig. 3e). To compare this circular embedding with the other tools, we quantified the circularity with circular ROC (CROC) ^57^. In 3 out of 4 stages, i.e., G1, mS and lS_G2, and on average, embedding by deDoc2.w had the highest CROC value (Fig. 3g). In eS, deDoc2.w came in second, along with GRiNCH, with comparable CROC. In G1 and mS, Higashi and GRiNCH came in second, along with deDoc.w, with comparable CROC, respectively. In general, the embedding by deDoc2, Higashi, GRiNCH, IS and scHiCluster showed a circular property, suggesting that TLDs do, indeed, carry information for cell cycle. Taken together, our assessment suggests that deDoc2 works well with experimental scHi-C data.

### deDoc2 can reveal hierarchical TLD and higher-level structure in single cells

To assess the ability of deDoc2 to identify hierarchical chromatin structure at single-cell level, we quantified the hierarchy of the domain structure with two metrics, modularity ^58^ and adjusted R^2^ (adjR^2, 34^). The average modularity and adjR^2^ for all TLDs in a sample were denoted as TLD-modularity and TLD-adjR^2^, respectively. For a given list of TLDs, a better-defined hierarchy will have larger TLD-modularity and larger TLD-adjR^2^.

With downsampled data at the sparse level of single-cell Hi-C, we found that deDoc2 identified hierarchical structure with larger TLD-modularity and larger TLD-adjR^2^ more stably when compared to two other hierarchy detectors, deDoc and SpectralTAD. The deDoc2-defined nested TLDs always yielded the highest WS and TLD-modularity (Fig. 4a, b). For TLD-adjR^2^, when the genomic distance is larger than 40kb, deDoc2 was always substantially higher than either deDoc or SpectralTAD (Fig 4c). Notably, when the genomic distance was smaller than 40kb, deDoc-defined nested TLDs yielded the highest TLD-adjR^2^. However, the length of TLDs and variation of adjR^2^s, as called by deDoc with genomic distance smaller than 40kb, were also much shorter and larger than those called by deDoc2, respectively (Fig. S5). Given the sparseness at the downsampling rates we tested (0.1%-0.025%), those small fragmented domains will be subject to severe stochastic fluctuation and thus unreliable. Therefore, deDoc2 can successfully define the hierarchy of a nested structure with ultra-sparse Hi-C data.

**Figure 4.**
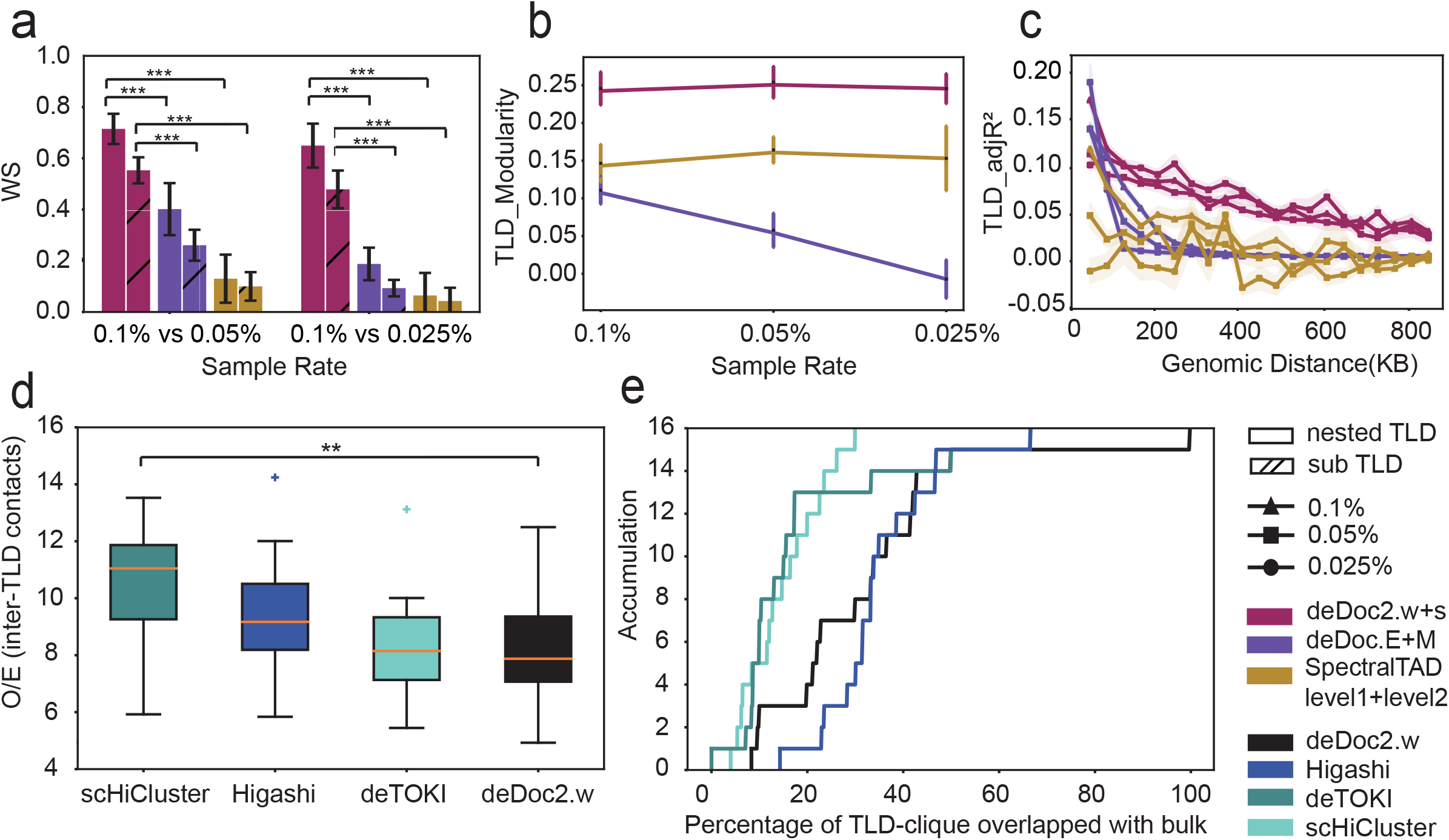
deDoc2 reveals hierarchical TLD with ultra-sparse Hi-C data at single-cell level. **a**. WS of nested TLDs and sub-TLDs identified by different tools at different downsample rate. Mann-Whitney U test ^∗∗^∗p<0.001, **p<0.01, *p<0.05 **b**. TLD_Modularity of nested TLDs identified by different tools at different downsample rate. **c**. TLD_adjR^2^ of nested TLDs identified by different tools at different downsample rate. **d**. Enrichment of inter-TLD contacts in TLD-cliques based on TLD results of different tools in Tan’s data. Mann-Whitney U test ^∗∗∗^p<0.001, **p<0.01, *p<0.05 **e**. Percentage of TLD-cliques overlapped with bulk Hi-C based on TLD results of different tools in Tan’s data.

Dedoc2 can also identify higher-level chromatin structure of TLDs, e.g., TLD-cliques. TAD-cliques comprise a higher level of chromatin structure than nested TADs found in bulk Hi-C data ^59^. So, we asked if such higher-level chromatin structure exists in single cells. Given the capacity of detecting TLDs in single cells, we expect well-defined single-cell TLD-cliques to meet the following requirements. First, they should have a clean TLD pattern, i.e., as few inter-TLD interactions as possible, given the significantly enriched inter-TLD interactions among TLDs in the clique. Second, they should be very accurate, considering consistency with bulk TAD-cliques in the cell population, cumulatively. We composed a pipeline to identify TLD-cliques at the single-cell level with predefined TLDs as input (Methods). We compared the cliques identified using TLDs called by deDoc2, scHiCluster, deTOKI and Higashi with Tan’s dataset ^53^. Within the cliques identified, deDoc2 showed the lowest O/E inter-TLD interactions (Methods), followed by deTOKI and Higashi (Fig. 4d). Thus, deDoc2-defined TLD-cliques are significantly cleaner than those of the other three tools in the single cells we examined. We further verified the accuracy of the identified TLD-cliques by overlapping them with bulk TLD-cliques called from the pooled single-cell Hi-C data. Altogether, deDoc2.w and Higashi identified the highest proportion of bulk TLD-cliques, higher than deTOKI and scHiCluster by more than 50% (Fig. 4e). These results show that deDoc2 can identify nested and higher-level structures in the hierarchy of chromatin architecture at the single-cell level.

### Hierarchical TLD structure is highly dynamic during cell cycle and among human brain prefrontal cortex cells

To investigate the dynamic hierarchical structure of chromosome architecture in single cells, we applied deDoc2 to two public single-cell Hi-C datasets, i.e., the cell cycle ^54^ and human brain prefrontal cortex cells ^55^. We found that the nested TLDs dramatically changed among cell types in both datasets. From G1 into early and middle S stage, both modularity and TLD_adjR^2^ were gradually decreased (Fig. 5 a-b), while from middle S to G2, they were increased and continued to drop for modularity and TLD_adjR^2^, respectively. Nevertheless, the absolute difference of TLD_adjR^2^ between middle S and G2 (0.145) was much weaker than the changes among earlier stages, i.e., 0.380 and 0.295 between G1 and both early S and middle S, respectively (Fig. 5b). Interestingly, we found that nested TLDs not only varied substantially among cell types in the human brain prefrontal cortex (Fig. 5 c-d), but they might also be used to classify cell types. Using hierarchical clustering, we found that single cells could be roughly classified into neural and non-neural categories by their modularity and TLD_adjR^2^ (Fig. 5e). Moreover, the two features could also largely pairwise distinguish the four categories of cells found in the prefrontal cortex (Glutamatergic, GABAergic, Non-Neuronal, and Non-Neural ^60^).

**Figure 5.**
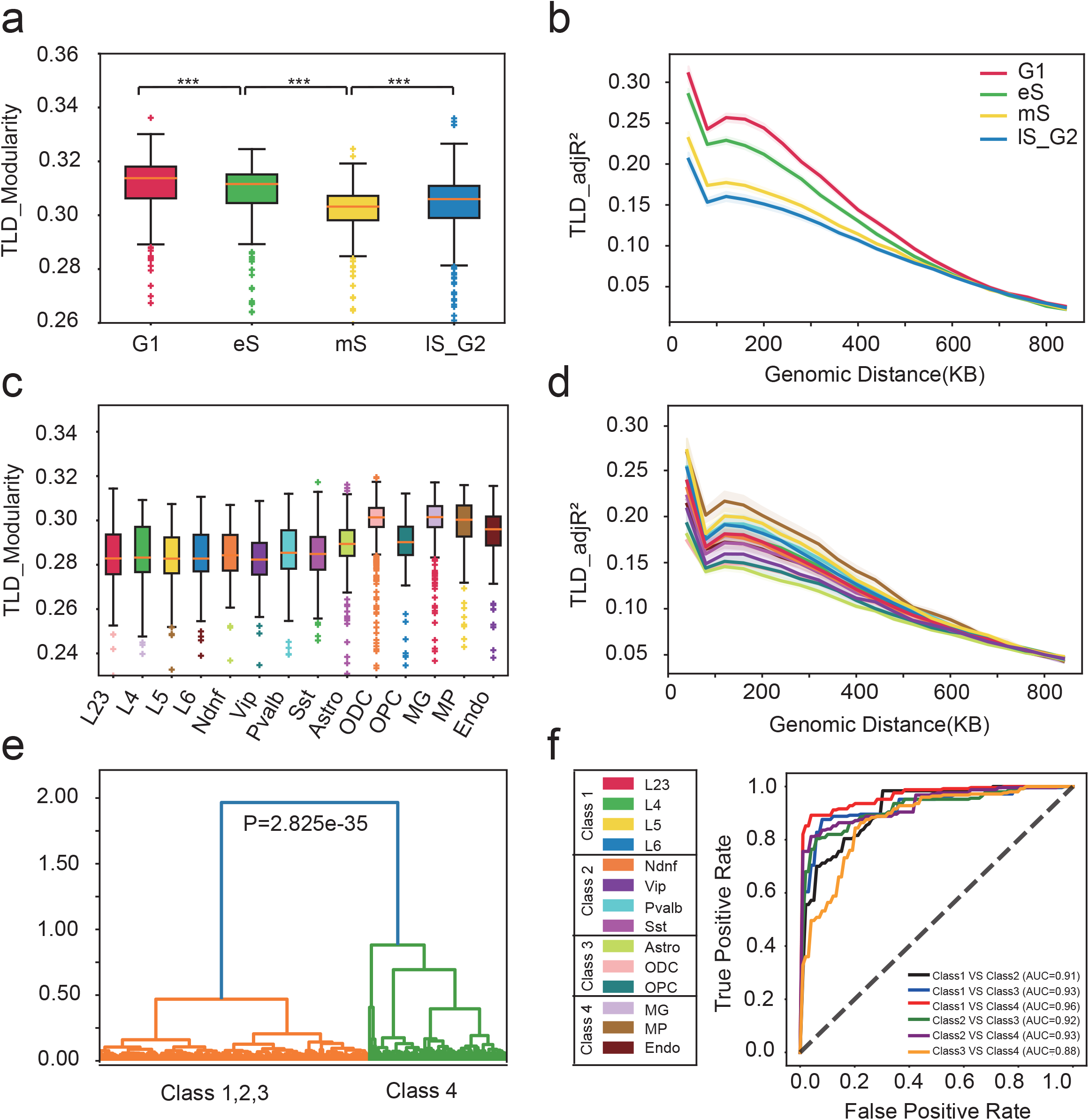
TLD structure is highly dynamic during cell cycle and among human brain prefrontal cortex cells. **a**. TLD_Modularity of TLDs predicted by deDoc2 with RWR imputation of mES cells in Nagano’s data (1137 cells). Mann-Whitney U test ^∗∗∗^p<0.001, ** p<0.01,* p<0.05 **b**. TLD_adjR^2^ of TLDs predicted by deDoc2 with RWR imputation of mES cells in Nagano’s data (1137 cells). **c**. TLD_Modularity of TLDs predicted by deDoc2 with RWR imputation of humanPFC cells in Lee’s data (4238 cells). **d**. TLD_adjR^2^ of TLDs predicted by deDoc2 with RWR imputation of humanPFC cells in Lee’s data (4238 cells). **e**. Hierarchical classification result of humanPFC cells in Lee’s data (30 cells randomly selected from each cell type, 420 cells in total). **f**. SVM classification result of humanPFC cells in Lee’s data (200 cells randomly selected from each cell type class, 800 cells in total).

Thus, our analysis may imply that the hierarchical structure of TLD nesting may carry key information for cell identity with concomitant functional implications. For example, cell type-specific genes were associated with cell type-specific nested TLD boundaries (Fig. 6a). We picked one representative cell type from each of the four cell classes, i.e., L23, Vip, Astro, and MG, and defined their cell type-specific highly expressed genes and nested TLD boundaries (Methods). We found that cell type-specific highly expressed genes were significantly enriched in cell type-specific nested TLD boundaries (Fig. 6a). To illustrate the functional implication for the hierarchical TLD, we cited apolipoprotein E (APOE). APOE is a key gene found to be associated with Alzheimer’s disease (AD)^61^. The APOE gene is highly expressed in Astro cells (Figure S6) and locates in an Astro cell-specific nested TLD boundary.

**Figure 6.**
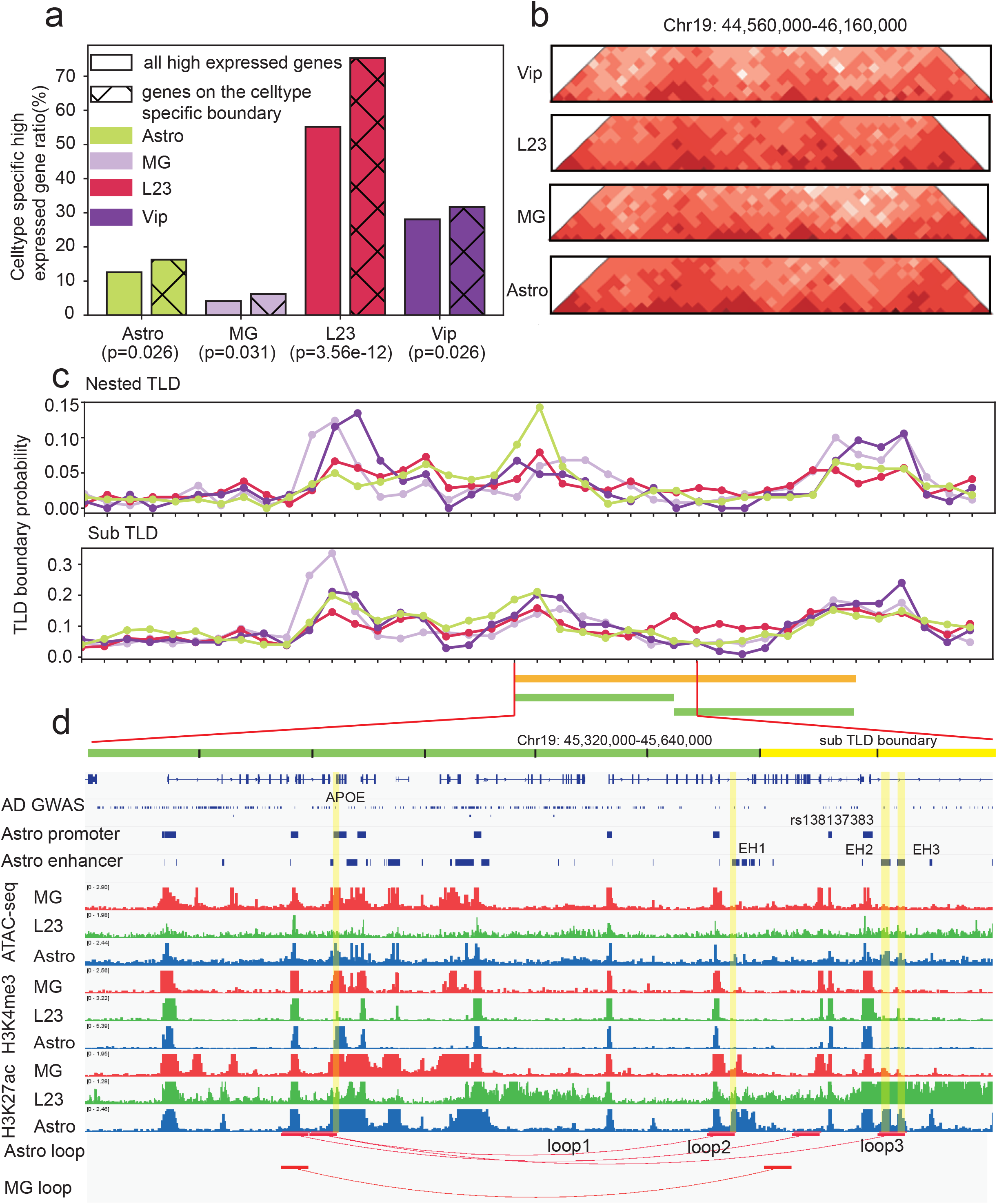
TLD structure related to cellular functionality. **a**. Cell type-specific highly expressed gene distribution of cell type-specific boundary and all genes. **b**. Pooled Hi-C matrix of four cell types at chr19:44,560,000-46,160,000. **c**. Nested TLD and sub-TLD boundary probability at chr19:44,560,000-46,160,000. **d**. Epigenetic data and loop identified by SnapHi-C around the APOE gene.

Two loops (loop1, loop3) in the locus link the APOE promoter to three active enhancers (EH1,EH2,EH3) identified by SnapHiC ^62^, of which the loop between EH2 and EH3 to APOE is Astro cell-specific. It is known that Astro cells are associated with AD^63-65^. EH2 and EH3 are also located in the Astro-specific sub-TLD boundary, and EH2 carries AD-associated GWAS SNPs (rs138137383) (Fig. 6b-d). Thus, Astro-specific EH2-APOE interaction between hierarchical TLD boundaries associated with the SNP towards AD is plausible. Needless to say, further detailed study is required to corroborate this speculation. Taken together, the hierarchical structure of chromosome architecture is a characteristic feature for single cells and may have profound effect on their cellular functionality.

## Discussion

In this work, we present a novel algorithm, deDoc2, to detect TLDs at single-cell level. Compared to its predecessor, deDoc, which reached local optimal of structure entropy, we implemented a dynamic programing algorithm to approach the global optimal of structure entropy in deDoc2. A recently published TAD caller, SuperTAD, also employed dynamic programing to approach global optimal of structural entropy. The deDoc2 and SuperTAD differ in that they minimize two- and high-dimensional structural entropy, respectively. SuperTAD requires several orders of magnitude higher CPU time and memory space to solve high-dimensional structural entropy (Supplementary Table 4). Moreover, high-dimensional entropy calculated from ultra-sparse scHi-C data may be less reliable. Therefore, SuperTAD may not be suitable for scHi-C data analysis. Compared to state-of-the-art tools, deDoc2 exhibits first-class performance on the metrics we assessed in the downsampled, simulated and experimental scHi-C data.

The deDoc2 distinguished itself most from peer predictors with several unique features. First, deDoc2 decodes TLD from single-cell Hi-C data with no need of data imputation, while most state-of-the-art TLD predictors do ^41^. In the ideal scenario, the imputation process does not introduce new information to the matrix regarding chromatin structure. However, the ultra-sparseness of scHi-C data made it plausible that certain artifacts may emerge from stochastic fluctuation. Thus, a method that does not involve data imputation may be less subject to the risk of false positives. Because the definition of structure entropy fully utilized the data in the whole contact matrix, deDoc2 achieved similar, even advanced, performance over imputation-based TLD predictors (SI and Fig. S2, S3).

Second, deDoc2 can reveal the structural hierarchy of TLDs. The hierarchical structure of TADs is a common property seen in bulk Hi-C data. However, to the best of our knowledge, it has been rarely discussed in the literature for single-cell Hi-C data, and the question may not be trivial. For example, whether the hierarchy seen in bulk Hi-C is an emerging property of heterogeneous cell population, or it does, indeed, exist in individual cells, remains largely unexplored. Our method showed a clear hierarchy in single-cell Hi-C data, and the brief examples we showed in the present work suggested that this hierarchy in single cells may carry fundamental biological functions. Thus, to further reveal the mechanisms of higher-level 3D genome folding, the dynamics and functions of hierarchical domain structure at the single-cell level will be essential. The experiments we showed here suggested that deDoc2 is a unique predictor in the spectrum of single-cell Hi-C data analytic tools.

Single-cell Hi-C technologies have been diverging into two distinct avenues, i.e., higher throughput in cell numbers with much sparser reads in each cell^54^ in contrast to higher data resolution in each cell with limited cells^53^. It remains challenging, even for deDoc2, to detect domain-like structure from a few thousands reads. Thus, breakthroughs of experimental single-cell Hi-C technologies and computational methods are both urgently needed in the field to enhance the throughput of single-cell 3D genome research. Moreover, to integrate with data from ligation-free methods, e.g., SPRITE ^66^ and ChiA-drop ^67^, even from imaging data, would be extremely helpful.

## Conclusions

We present a newly developed single-cell Hi-C data analysis tool termed as deDoc2. Developed from structural information theory, deDoc2 not only outperformed state-of-the-art tools for TLD detection, but it also has the unique feature of detecting the hierarchy of chromosome domain structure in single cells. Finally, we demonstrated that higher-level single-cell 3D chromatin structure dramatically changes during cell cycle and among cell types and that this dynamism is tightly associated with gene expression changes, implying its functionality.

## Methods

### Structural information theory

Structural information theory ^45^measures and decodes information embedded in a system consisting of many bodies together with interactions among the individuals of the several bodies^1^. Hi-C data can be interpreted as a weighted graph, and the TLD detection problem can be viewed as a graph partitioning problem. As in deDoc, we seek to detect TLDs by structural information theory.

A fundamental concept in structural information theory is the notion of encoding a tree, i.e., a connected graph with no cycles. In fact, encoding a tree is both mathematical model and data structure of the hierarchical abstracting of a graph.

#### Encoding tree

Structural information theory measures the information embedded in a graph that is decoded by an encoding tree. An encoding tree *T* of a graph *G*(*V,E*) is defined as follows:

1. For every tree node *α* ∈ *T*, a vertices set *T*_*α*_ ∈ *V* is associated with it, and the immediate successors of *α* are labeled by *α*^^^ ⟨0⟩, *α*^^^ ⟨1⟩ …, *α*^^^ ⟨*k*⟩ We say that *α* is the codeword of the set *T*_*α*_, and that *T*_*α*_ is the marker of *α*.
2. The root of *T* is an empty string *λ*, associated with the vertices set *V*, written as *T*_*α*_ = *V*.
3. For every *α* ∈ *T*, if *β*_0_, *β*_1_, …, *β*_*k*_ are all immediate successors of *α* in *T*, then 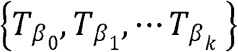 is a partitioning of *T*_*λ*_.
4. For every leaf node *λ* ∈ *T, T*_*λ*_ contains only one vertex of *V*.

The structural entropy of graph *G* given by an encoding tree *T* is defined as

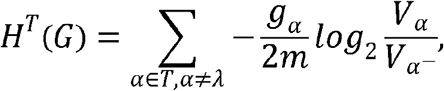

where *α* is a tree node of the encoding tree *T, λ* is the root of *T, α*^−^ is the immediate predecessor of *α, g*_*α*_ is the cut of *α*, i.e., the number of edges between the vertices in and not in vertices *T*_*α*_, *V*_*α*_ is the volume of *α*, i.e., the sum of degrees of all vertices in *T*_*α*_, and *m* is the sum of the edges, *sum*(*E*). The structural entropy of *G* is *min*_*T*_ {*H*^*T*^(*G*)}, and the *k*-dimensional structural entropy of *G* is *min*_*T*_ (*H*^*T*^(*G*)}, where *T* has height restriction of at most *k*.

#### Decoding information

Given a graph *G* and an encoding tree *T* of *G*,

i. the metric *H*^1^(*G*), i.e., the one-dimensional structural entropy of *G*, is the total amount of uncertainty that is embedded in *G* and
ii. the metric *H*^*T*^(*G*) is the amount of uncertainty that is embedded in the system obtained from *G* by the hierarchical abstracting given by *T*.

Therefore, the information decoded by the hierarchical abstracting *T* of encoding tree *T*, referred to as the decoding information of encoding tree *T* from *G*, is

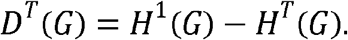

The decoding information of *G* is *max*_*T*_ {*D*^*T*^(*G*)}, and the *k* -dimensional decoding information of *G* is *max*_*T*_ {*D*^*T*^(*G*)}, where *T* has height restriction of at most *k. D*^*T*^(*G*), is the metric of the information that is decoded by the strategy of hierarchical abstracting given by encoding tree *T* from graph *G*.

### deDoc2

Aiming to seek the optimal partition of genome, deDoc2 employed a dynamic programming algorithm to minimize two-dimensional structural entropy. The deDoc2 package has 3 components. DeDoc2.w and deDoc2.s predict the global and local partition of genome, respectively, and deDoc2.binsize determines the optimal binsize with normalized decoding information.

#### deDoc2.w: minimize two-dimensional structural entropy by dynamic programming algorithm

Given a graph *G*, an encoding tree *T* with height 2 of *G* is equivalent to a vertex partitioning of *G*.

DeDoc2.w predicts the optimal partition *P* = {*X*_1_, *X*_2_, …, *X*_*L*_}, where each *X*_*i*_ is a collection of continuous bins representing a TLD in the genome with minimal two-dimensional structural entropy. By the definition of structural entropy of a graph under an encoding tree, the two-dimensional entropy of *G* given by *P* can be expressed as

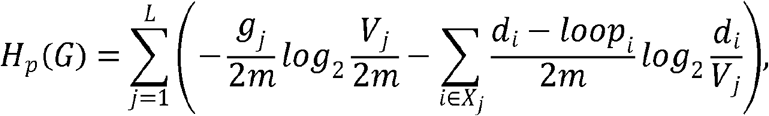

where *d*_*i*_ is the degree of node *i, loop*_*i*_ is the weight of self-loop edge of node *i*, i.e., the *i*-th value of diagonal, *g*_*j*_ is the cut of TLD *j, V*_*j*_ is the volume of TLD *j*, and *L* is the number of partitions. When predicting TLDs of a chromosome, if we put *k* + 1 bins from bin *i* to bin *i* + *k* (bin *i* and bin *i*+*k* included) in a TLD, we define the two-dimensional structural entropy of this TLD as

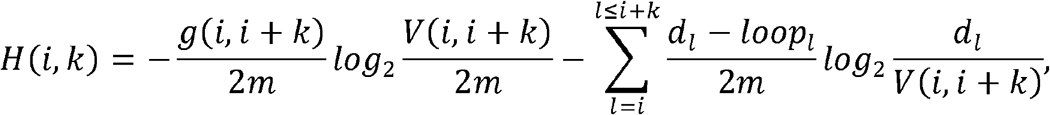

where *g*(*i, i* + *k*) is the cut of this TLD, and *V*(*i, i* + *k*) is its volume. The recurrence relation can be written as

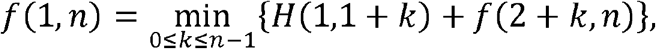

where *f*(1, *n*) is the two-dimensional structural entropy of a graph that includes nodes from 1 to *n*. To speed up the algorithm, we limit the length of TLDs within 10Mb, i.e., 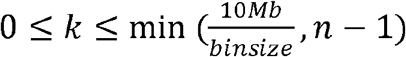, in the recurrence relation equation above.

There are n possible values for k to choose and n possible *H*(1,1 + *k*) terms, and *H*(1,1 + *k*) can be calculated in *o*(1) time, so the time complexity of deDoc2.w is written as *o*(*n*^2^). With the limitation of TLD length within 10Mb, the possible value of k becomes 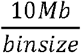, and the time complexity becomes 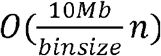, which is linear to the length of chromosome.

To accommodate isolated vertices, which are usually seen in sparse Hi-C data, we add a self-loop value 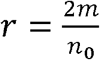, where *n*_0_ is the number of non-isolated vertices in the matrix, to increase the aversion of these nodes to form communities (see SI).

#### deDoc2.s: minimize two-dimensional structural entropy in sliding windows by dynamic programming algorithm

When two-dimensional structural entropy minimization is applied to a smaller matrix, the TLD structure will be smaller. We use a sliding window to split the Hi-C contact matrix and apply deDoc2.w to the split matrix to get smaller TLDs and form a hierarchical structure with TLDs, which, in turn, is predicted by deDoc2.w. We set default window size to be 10 Mb, which means 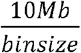 bins in the matrix. The first sliding window starts from the first bin and ends at 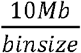 bin. After TLDs in the first window are predicted by deDoc2.s, the window slides to the first bin of the last TLD predicted to keep potential TLDs from being cut by the sliding window. Repeating this to the end of the contact matrix, we get the predicted TLDs.

The time used for predicting TLDs in one sliding window is 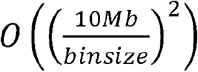, where 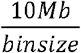 is the number of nodes in a sliding window, and the time complexity of deDoc2.s is written as 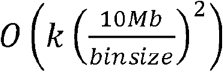, where *k* is the number of sliding windows.

### deDoc2.binsize: Normalized decoding information for binsize determination

Decoding information of a graph measures the maximum amount of information that can be eliminated by an encoding tree such that a graph with larger decoding information implies a better community structure. In particular, in the case of Hi-C data, larger decoding information implies a better TLD structure. Our aim is to choose a proper binsize with which to build a graph that best shows the TLD structure in Hi-C data. We only calculated 2D structural entropy in this paper, so we chose *D*^2^(*G*) as the decoding information for binsize determination. For simplicity, we used the encoding tree obtained by deDoc2.w to calculate the decoding information since deDoc2.w can get the optimal 2D encoding tree that most minimizes the structural entropy of the graph. Because decoding information is not comparable between binsizes, we searched for a way to normalize the decoding information ^68^. To do this, we normalized the residual entropy, i.e., the decoding information of a partition, by dividing the 1D structural entropy of the graph. Here we did the same normalization and define the normalized decoding information (NDI) as

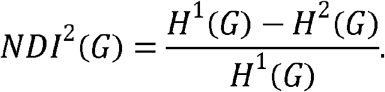

We propose to choose a binsize which maximizes the NDI of the Hi-C contact matrix. We chose a binsize so that its associated Hi-C graph has the minimum normalized SE (nSE) among all stable binsizes in ^33^. Comparing NDI and nSE, we find that if nSE is minimum, the NDI is maximum among binsizes, so NDI and nSE are consistent with each other. However, it is worth noting that 2D structural entropy is calculated by deDoc(E) in nSE, while it is calculated by deDoc2.w in NDI; thus, the resulting structural entropy can be slightly different.

### Imputation methods

#### Random walk with restart (RWR) imputation

Random walk with restart (RWR) provides a good relevance score between two nodes in a weighted graph, and it has been used in numerous settings ^69^. For a sparse Hi-C matrix *M* to be imputed, we normalize it into a matrix *C* by sqrtVC normalization ^50^, which ensures that the resulting RWR imputation can converge and keep *C* symmetric. The imputed matrix *Q*_*t*_ is calculated recursively as

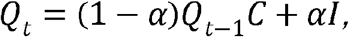

where *Q*_0_ =*I*, and *α* is the restart probability. The random walk stops when || *Q*_*t*_ − *Q*_*t*−1_ ||_2_ ≥ 10^−2^, which means that the process has converged.

### Adding arbitrary noise

Noise was introduced to the Hi-C matrix as follows ^70^. For a given Hi-C matrix (*N*×*N*), a proportion (*P*_*n*_) of entries were randomly selected with replacement and a constant of one added. As the selection was with replacement, the numbers added into the matrix follow Binomial distribution *B* (*n,p*), where *n* is the number of entries to select and *p* is the *1/N*^*2*^. The *P*_*n*_ was set to be 10%, 20%, and 40% of the number of contacts in the original downsampled Hi-C matrices (0.025%) to form three noise rates. Considering that the contact probability of Hi-C matrices decreases dramatically as the genomic distance increases, we calculated the contact probability of each secondary diagonal using the method in Boost-HiC ^71^ and randomly selected entries with replacement of each secondary diagonal separately according to its contact probability. To avoid adding noise to unmappable regions, we only considered the rows/columns with at least one nonzero entry in the downsampled Hi-C matrices.

### The simulation of scHi-C data

We simulated scHi-C data according to our previous work^42^. Briefly, a 3D model was simulated using IMP ^72^, and we simulated the reference and scHi-C data as follows^73^. For reference Hi-C, reads were sampled from any two genome loci *i* and *j* with the weight defined as *Weight(i, j) = 1/distance(i, j)*. For scHi-C data, reads from any two genome loci *i* and *j* were sampled with the weight defined as *Weight(i, j) = D-distance(i, j)*,where D is a threshold. Only genome loci having Euclidean distance less than D were considered to be contacting. The simulation generated data with 40kb resolution, and the Hi-C contact matrix was used for actual TLD detection (see details of simulation from the original paper^42^).

### Execution of TAD predictors

All test TLD predictors were run with default settings, except for Higashi^49^, which we set to be *nbr=4* for multicell datasets and *nbr=0* for downsampled and simulated Hi-C data. We also set *embedding_epoch=1* and *no_nbr_epoch=20* for downsampled and simulated Hi-C data to save time, and, as suggested by the authors, the TADs of bulk Hi-C data were called by the function *call_tads* in Higashi codes. CPU times can be found in the Supplemental Table 5.

### Adjusted Mutual Information AMI (T, K)

Mutual information MI (T, K) was defined as

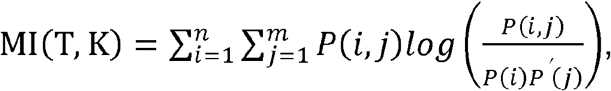

where

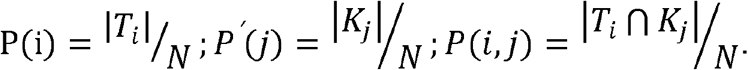

Then, the adjusted mutual information AMI (T, K) was defined as

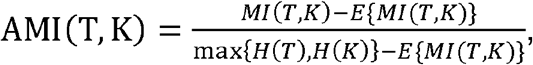

where *H* denotes the Shannon entropy, and *E* denotes expectation. AMI was calculated by the function adjusted_mutual_info_score in the Python module sklearn.metrics. In real calculation, all predicted TLDs and intermediate windows of TLDs are included in *T* and *K*.

### Weighted Similarity WS(T, K)^33^

The weighted similarity WS(T,K) was defined as

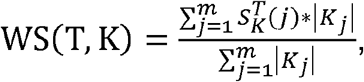

where

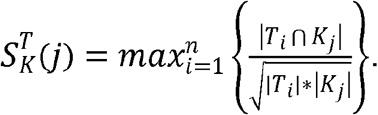

Because WS is an asymmetric index for similarity, we always put the predicted TLDs from raw data in *T* and the TLDs from downsampled data in *K*, while the intermediate windows of the domains were not included in either *T* or *K*.

### The enrichment of structural proteins and histone marks around TLD boundaries

ChIP-seq peaks data of structural proteins and histone marks for Nagano’s dataset were downloaded from Yue, Feng, et al. ^74^. ChIP-seq peaks data for all other datasets were downloaded from ENCODE (www.encodeproject.org) ^75^. To quantify the enrichment of ChIP-seq peaks around TLD boundaries, we calculated the fold change between peaks found at TLD boundaries and those at adjacent flanking regions ^76^. The boundary regions were defined as TLD boundaries with two flanking bins, and flanking regions 100kb and 500kb away from the TLD boundaries were defined as background. The fold change of peaks is calculated as 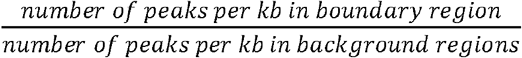.

The definition of TLD boundaries. For predictions by score-based detectors, e.g., IS, deTOKI, scHiCluster and Higashi, boundaries were defined as the predicted bins, and the fold changes of ChIP-seq were calculated at the midpoint of the bin. For predictions by network-based TLD detectors, e.g., deDoc, deDoc2, SpectralTAD and GRiNCH, boundaries were defined as the edges between two predicted bins, and the fold changes were calculated at the edges.

### Structural entropy index

We interpret Hi-C contact matrix *M* as a graph and used TLDs predicted by different algorithms to build an encoding tree *T* with height two. We then calculated the structural entropy of *M* given the encoding tree *T* (two-dimensional structural entropy). The vertices in the gap regions were put directly on the root *λ* of *T* since these vertices do not belong to any communities.

### Modularity index

For a Hi-C contact matrix *M*, we replaced the diagonal elements with zero, and the remaining elements were log2 processed. Modularity ^77^ was defined as

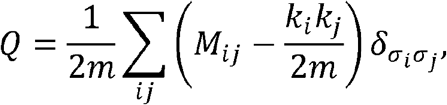

where *i* and *j* are bin indices of *M, k*_*i*_ and *k*_*j*_ are the degrees of the vertices, and 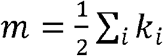 is the total number of edges in the network. The value of Kronecker delta 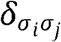 equals 1 if *σ*_*i*_ = *σ*_*j*_, where the label *σ*_*i*_ denotes the community label of bin *i*. For the gap region recognized by algorithms, we take each single locus of the vertices in these regions as a community.

### Embedding and clustering of single cells

The visualization of embedded single cells of TLDs predicted by deDoc2 utilized the result of deDoc2 with RWR imputation, i.e., 4238 cells and 1171 cells used in Lee’s and Nagano’s datasets, respectively. The AMI and CROC to quantify the embedding used 400 and 560 randomly selected cells in Lee’s and Nagano’s datasets, respectively.

### Visualization of embedded single cells

Following the visualization method, as described in Zhou and colleagues ^41^,we performed two-dimensional embedding using TLD predictions as text-document input on Lee’s dataset. For Nagano’s dataset, we omitted one step in the above visualization method, i.e., harmonypy ^78^, as notable batch effect was spotted which may negatively affect the dimension-reducing results. Last, we applied t-distributed stochastic neighbor embedding (T-SNE) to the dimensionality-reduced matrix to get a two-dimensional map of the data for visualization by the Python module MulticoreT-SNE.

### CROC

We used the visualization result described above to calculate circular ROC ^57^ (CROC) to evaluate the quality of embedding from TLD boundaries of a cell-cycle dataset. For a multiclass dataset, we calculated CROC for one class at a time by taking all remaining classes as negative points. For a circular two-dimensional cell embedding, we get a unique angle *θ* for each cell. We assumed that the angles of a class follow a von Mises distribution and calculated the mean angle *θ* ^*^ as the true positive label of the class; then we took the difference between *θ*^*\**^ and every angle *θ*_*i*_ of a data point as the predicted score of this data point. The ROC calculation was then performed by sklearn.metrics.roc_auc_score. To evaluate the embedding of all cell types, we calculated average CROC (ACROC) across these types.

### Evaluation of the embedding of non-circular datasets

To evaluate the embedding of non-circular datasets, we clustered the embedding and calculated the AMI score ^52^, After the first dimensionality reduction as we did in the visualization step, we applied k-means clustering on the first 15 dimensions of the dimensionality-reduced matrix and calculated the AMI score by the Python module sklearn.metrics.adjusted_mutual_info_score.

### Hierarchical TLD assessment

We regard each nested TLD as an independent matrix and calculate modularity according to the division of the subTLDs. TLD modularity is the average modularity of all nested TLDs in the cell.

TLD-adjR^2^ was adopted from An, L. et al^34^, and it can measure the proportion of Hi-C signal variation explained by TAD calls. For any given genomic region and given genome distance, the TLD-adjR^2^ is defined as

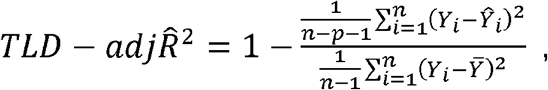

where *Y*_*i*_ denotes the contact number of the *i-*th bin, *n* denotes the number of bins at the same genomic distance as this bin, and *p* denotes the number of subTLDs larger than, or equal to, the genomic distance. For bins within subTLD, *Ŷ*_*i*_ denotes the average contact frequency at a given genomic distance within that subTLD. For those bins not in any subTLDs, *Ŷ*_*i*_ is the average of contact frequency in the gap region at that genomic distance, and 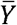 denotes the overall mean contact frequency across all bins at a given genomic distance.

### Identification of TLD cliques

The pipeline for TLD clique identification was adopted from Paulsen, J. et al ^59^, we only modified the TAD with TLD during the whole process.

### The enrichment of inter-TLD contacts in TLD-cliques

We adopted three rules from Paulsen, J. et al ^59^: (i) the permuted clique was on the same chromosome as the observed clique; (ii) sizes of TLDs in a clique were kept; (iii) genomic distance between consecutive TLDs in the clique was maintained. The start position of a TLD clique was randomly positioned on the same chromosome and TLD size and inter-TLD distances were shuffled to construct a permuted TLD clique. In total, 250 permutations were carried out and the expected inter-TLD contacts is the mean inter-TLD contacts of the 250 permuted cliques. The enrichment of inter-TLD contacts in TLD-cliques is calculated as (observed inter-TLD contacts)/ (expected inter-TLD contacts).

### Cell type-specific highly expressed genes and nested TLD boundaries

To get cell type-specific highly expressed genes, we first get the single-cell expression data of Astro, MG, L23 and Vip from the single-cell RNA-seq dataset of brain (https://portal.brain-map.org/atlases-and-data/rnaseq/human-multiple-cortical-areas-smart-seq). Then, if the expression level of gene A in certain cell type was higher than that of the other three cell types, gene A was regarded as the corresponding high-expression gene of this cell type.

As for cell type-specific nested TLD boundaries, the frequency of occurrence of a boundary in each of the four cell types was first calculated. If the frequency of occurrence of a boundary was higher than the mean plus three standard deviations, it was called a high frequency boundary. If the 3 bins before and after this position did not have a high frequency boundary in other cell types, then that boundary was set as the boundary specific to this cell type.

## Declarations

### Availability of data and materials

The source code can be freely accessed through github at https://github.com/zengguangjie/deDoc2. All simulated and experimental data used in this study are summarized in Supplementary Table 6. No Hi-C data were normalized.

### Funding

This work was partially supported by Special Investigation on Science and Technology Basic Resources of the MOST, China (2019FY100102), the Strategic Priority Research Program of the Chinese Academy of Sciences, China (XDA24020307), the National Key R&D Program of China (2018YFC2000400), Beijing Natural Science Foundation (Z200021), and by the National Nature Science Foundation of China (61932002, 61772503, 31871331, 91940304).

### Authors’ contributions

AL, GZ and ZZ conceived this project. GZ and HW performed the experiments. GZ, HW and XL analyzed data. AL, GZ, HW and ZZ prepared the manuscript. All authors read and approved the final manuscript.

### Competing interests

The authors declare that they have no competing interests.

### Ethics approval and consent to participate

Not applicable.

### Consent for publication

Not applicable.

## Acknowledgements

Some experiments in this paper were carried out on the High Performance Computing Platform of Beihang University. We thank Mr. David Martin for English language editorial services.

## Footnotes

1 Angsheng Li, Mathematical Principles of Information World: Calculus of Information

## Notes

### Competing Interest Statement

The authors have declared no competing interest.

