## Supplementary information for "DeDoc2 identifies and characterizes the hierarchy and dynamics of chromatin TAD-like domains in the single cells"

**Determining binsize using normalized structural entropy**

Here we propose to use structure information to determine binsize for a given Hi-C contact matrix. The scale of a graph can vary significantly when binsize changes. Since structure entropy is not comparable between binsizes, we seek to properly normalize structure entropy. Previously, we proposed to normalize structure entropy by dividing 1D structural entropy of the graph^1^ ^2^. We calculated the normalized structure entropy (NDI) of 23 chromosomes of GM12878 cell 14 from Tan’s data with different binsizes ^3^ (Fig. S1a), and found that most (20 out of 23) chromosomes had a bell-shaped distribution of NDI over binsize. Thus, the binsize with maximal NDI was chosen to build up the graph in Hi-C data. However, sometimes the highest NDI was reached at multiple binsizes. For example, in chromosome 18, all NDIs reached maximal with binsize=85kb, 135kb, and 170kb (Fig. S1b, c and d). TLDs can be detected in all binsizes, but TLDs tend to be longer when binsize=170kb. Smaller binsize can help retain subtle TLD structure, while larger binsize, albeit within a reasonable range, can help show TLD structure in Hi-C data. Thus, we propose to first choose an acceptable binsize range for TLD analysis and then choose the binsize that reaches maximum NDI.

**Improvement of deDoc2 with data imputation**

Data imputation is a commonly used strategy when handling single-cell data, e.g., scHiCluster and Higashi. We examined the improvement in performance for TLD predictors with imputation methods adapted from scHiCluster ^4^Higashi ^5^ and RWR, hereinafter denoted as scHiCluster, Higashi and RWR, respectively (see Methods, Fig S2., Fig. S3).

Using sample rate at 0.025% with the dataset of Rao et al. as an example ^6^, we examined the performance of 6 TAD predictors. First, we found that no data imputation methods we tested were universal for TAD detection. For example, Higashi can remarkably improve the performance of SpectralTAD, but it introduces a negative effect on most other TAD predictors (Fig. S2a-S2b). We found that the data imputation methods RWR and scHiCluster could slightly improve the performance of deDoc2, as measured by AMI and WS (P < 0.001, paired sample *t*-test) (Fig. S2a-S2b). However, such improvement was barely seen in the enrichment of structural protein and histone marks in the boundaries (Fig. S3a). As for other TLD predictors, we found that RWR and scHiCluster imputation could improve AMI, WS and the enrichment of CTCF. No evident improvement could be seen for all other predictors under different imputation methods.

We then tested the effect of data imputation for simulated scHi-C data ^7^ (Fig. S2c-S2d, S3b-S3c). We noted slight improvement of WS between simulated scHi-C TLDs and simulated bulk TLDs for deDoc2, IS and deDoc under RWR and scHiCluster imputation for threshold=500 (Fig. S3b-S3c) and threshold=750 (Fig.S2c-S2d).

We also tested the effect of data imputation for experimental scHi-C data^3^. While the improvement of modularity under different imputation methods can be seen for most predictors, the improvement of TF binding enrichment is not remarkable (Fig. S2e, S3d). We also tested the improvement of TF binding enrichment and CROC of TLD embedding for Nagano’s data ^8^, and the improvement of AMI of TLD embedding for Lee’s data ^9^. No evident improvement can be seen for predictors for either one (Fig. S2f, S3e, S3f). However, we did find that Higashi imputation showed improvement for most predictors for Nagano’s data (Fig. S2f, S3e), and scHiCluster imputation showed improvement for most predictors for Lee’s data (Fig. S3f).

Together, at least for the data and tools we tested, the improvement from data imputation was minor and highly method-dependent. Among all these imputation methods and predictors, we found that RWR may be the best choice for deDoc2; in most cases we tested, it benefited deDoc2 with little defect.

Next, we examined improvement of the identification for nested and TAD-cliques by data imputation in Tan’s dataset. For nested TADs, the modularity of native deDoc2 was larger than that of SpectralTAD with data imputation (Fig. S2g). For TLD_adjR^2^, however, data imputation, in general, had a negative effect on deDoc2 (Fig.S2h), i.e., native deDoc2-generated TLDs had the highest TLD_adjR^2^ for all genomic distances. SpectralTAD-generated TLDs did, indeed, have a slightly higher TLD_adjR^2^ when the data were imputed by Higashi. However, this improved TLD_adjR^2^ remains smaller than that of native deDoc2. As for TLD-cliques, both deDoc2.w and deTOKI are more or less improved by certain data imputation methods; however, such improvements were minor, as well. For example, slightly more and clear TLD-cliques overlapped with bulk could be identified with RWR for deDoc2.w and with scHiCluster for deTOKI (Fig. S2i and j). Once again, the improved number of TLD-cliques remains smaller, even after data imputation for the other predictors. Altogether, our data suggest that data imputation leads to no substantial improvement in TLD prediction with single-cell Hi-C data.

**Performance of deDoc2 in bulk Hi-C data**

We assessed the performance of deDoc2 in bulk Hi-C data by comparing it with IS and deDoc using 6 cell types from the dataset of Rao et al. (Fig. S7). deDoc is a hierarchical TAD detector designed for bulk Hi-C data containing two predictors, deDoc(E) and deDoc(M). In bulk Hi-C data, we assessed both deDoc(E) and deDoc(M). In 6 cell types and 5 algorithms, all predicted a sufficient number of TADs, and the lengths of predicted TADs were all reasonable (Fig. S7a, S7b). deDoc2.w and deDoc(E) predicted TADs in comparable number and length, and deDoc2.s and IS predicted TADs in comparable number and length, while deDoc(M) predicted more TADs with smaller TAD length. We calculated fold change between CHiP-seq peaks found at TAD boundaries and those at adjacent flanking regions to evaluate the enrichment of structural protein and histones (CTCF, H3K4me3 and H3K36me3) on TAD boundaries. All algorithms have some degree of enrichment on the TAD boundaries they predicted (Fig. S7c). The enrichment of CTCF on TAD boundaries predicted by deDoc2.s and deDoc2.w ranked first and second, respectively, and the enrichment of H3K4me3 and H3K36me3 on TAD boundaries predicted by deDoc2.s and deDoc2.w ranked second and first, respectively, for most cell types. This indicates that deDoc2 performed best among the three algorithms in predicting TADs. We calculated modularity and structural entropy on Hi-C contact matrix of 6 cell types and calculated contact density on GM12878 Hi-C contact matrix. We found that TADs predicted by deDoc2.w and deDoc(E) had the highest modularity, lowest structural entropy and highest contact density and that these indices were comparable to each other (Fig. S7d-f). Modularity, structural entropy and contact density of TADs predicted by deDoc2.s and IS were also comparable.

**Difference between deDoc2 and SuperTAD**

Both deDoc2 and SuperTAD ^10^ use dynamic programming algorithm to minimize structural entropy of the Hi-C contact matrix to predict domains, but they differ in several respects. First, deDoc2 minimizes two-dimensional structural entropy and generates partitions as results, while SuperTAD minimizes high-dimensional structural entropy and generates hierarchical TADs as results directly. Multilevel hierarchical TAD structure is the advantage of SuperTAD; however, it has greater time and space complexity (Fig S8a). The time complexity of SuperTAD, both with and without height restriction of the coding tree, is $O\left( k^{2}n^{3} \right)$ and $O\left( n^{4}k^{2}h \right)$ , respectively, while the time complexity of deDoc2 is about $O\left( n^{2} \right)$, where $n$ is the number of bins, and $k$ is the number of encoding tree leaves. The definition of encoding tree leaves of SuperTAD is different from that of deDoc2. Tree leaves of SuperTAD indicate the lowest level of communities, which is comparable to the partition of deDoc2. To compare the performance of deDoc2 and SuperTAD on sparse Hi-C data, we tested these two algorithms on simulated scHi-C data, downsampled Hi-C data and experimental scHi-C data. Since SuperTAD outputs hierarchical TADs, we used the smallest TADs in the hierarchical structure in the following analysis to compare with deDoc2.

We first tested SuperTAD and its variant SuperTAD(F) (SuperTAD with filter), SuperTAD(2) (SuperTAD with height constraint $h=2$), and deDoc2 on simulated single-cell and bulk Hi-C data (see Methods). These Hi-C data are simulations of 5Mb chromosomes with binsize=40kb, which means that the Hi-C graph contains 125 vertices. We found that SuperTAD could predict TLDs of these short Hi-C fragments using time comparable to that of deDoc2. SuperTAD(2) spent about 1000-fold more CPU time than deDoc2 and SuperTAD (Fig. S8b). To evaluate the accuracy of predictions, we calculated the similarity between TLDs of simulated single-cell Hi-C and TLDs of simulated bulk Hi-C, and we found that the similarity of TLDs predicted by deDoc2 was clearly higher than that of TLDs predicted by SuperTAD, SuperTAD(F) and SuperTAD(2). For D=750, D=1000, and D=500, the similarity of TLDs predicted by deDoc2 was higher than that of SuperTAD(2) (Fig. S8c-d).

Second, we tested SuperTAD, SuperTAD(F) and deDoc2 on down-sampled IMR90 Hi-C data from Rao’s dataset at sampling rate 0.025% and on experimental GM12878 scHi-C data from Tan’s dataset. In the 9 shortest chromosomes (chr14 to chr22), we found that SuperTAD spent about 10000-fold more time than deDoc2 to predict TLDs (Fig. S8a and e) and that TLD boundaries of downsampled Hi-C data predicted by SuperTAD and SuperTAD(F) had only slightly higher enrichment than those of deDoc2.w and deDoc2.s. The enrichment of TLD boundaries of experimental single-cell Hi-C data predicted by deDoc2.w was comparable to that of SuperTAD and SuperTAD(F).

**Treatment of isolated vertices**

Taking Hi-C contact matrix as a graph, isolated vertices are bins that have no interactions with other bins. These isolated vertices could be bins in unmappable regions or bins with no reads mapped. Isolated vertices were either dropped or put into a TLD according to their genomic context. All bins in unmappable regions were simply dropped (Fig. S9a). The remaining isolated vertices will also be dropped in cases where only a few Hi-C reads connect the flanking region of bins (Fig. S9b). We add a self-loop value$r=\frac{2m}{n_{0}}$, where $n_{0}$ is the number of non-isolated vertices in the matrix, to increase the aversion of these nodes to form communities, which helps deDoc2 to determine whether drop isolated vertices or not.

**Figure S1.** deDoc2.binSize: Determine binsize using normalized structure entropy. Example Hi-C data was taken from Tan et al.^3^ GM12878 cell 14 chr18:18-35Mb.

a. NDI of 23 chromosomes of GM12878 cell 14 from Tan’s dataset in different binsizes.
b. NDI of chr18 of GM12878 cell 14 from Tan’s dataset in different binsizes. NDI reaches highest at binsize=170kb, while NDIs at binsize=85kb and binsize=135kb are also high.
c. Result of deDoc2 at binsize=85kb of chr18 of GM12878 cell 14 from Tan’s data.
d. Result of deDoc2 at binsize=170kb of chr18 of GM12878 cell 14 from Tan’s data.

**Figure S2.** Improvement of performance with data imputation.

a. Improvement of AMI between TLDs from downsampled Hi-C at downsample rate of 0.025% in Rao’s ^6^data.
b. Improvement of WS between TLDs from downsampled Hi-C at downsample rate of 0.025% in Rao’s data.
c. Improvement of AMI between TLDs from simulated single-cell Hi-C at threshold 750 and bulk Hi-C.
d. Improvement of WS between TLDs from simulated single-cell Hi-C at threshold 750 and bulk Hi-C.
e. Improvement of the modularity of predicted TLDs from GM12878 experimental single-cell Hi-C from Tan’s data.
f. Improvement of CROCs of embedding of mES cells in Nagano’s data (400 cells).
g. Improvement of TLD_modularity of predicted nested TLDs.
h. Improvement of TLD_adjR2 of predicted nested TLDs.
i. Improvement of enrichment of inter-TLD contacts in TLD-cliques of predicted TLDs.
j. Improvement of credible TLD-clique (size>5) percent of predicted TLDs.

\# means P < 0.05, * means P < 0.001, paired *t*-test.

**Figure S3.** Improvement of performance with data imputation.

a. Improvement of ChIP-seq peak signal (CTCF, H3K4me3 and H3K36me3) enrichment at TLD boundaries. TLDs are called from downsampled Hi-C at downsample rate of 0.025% in Rao’s data.
b. Improvement of AMI between TLDs from simulated single-cell Hi-C at threshold 500 and bulk Hi-C.
c. Improvement of WS between TLDs from simulated single-cell Hi-C at threshold 500 and bulk Hi-C.
d. Improvement of ChIP-seq peak signal (CTCF, H3K4me3 and H3K36me3) enrichment at TLD boundaries. TLDs are called from experimental GM12878 single-cell Hi-C in Tan’s data.
e. Improvement of ChIP-seq peak signal (CTCF, H3K4me3 and H3K36me3) enrichment at TLD boundaries. TLDs are called from experimental GM12878 single-cell Hi-C in Nagano’s ^8^ata.
f. Improvement of AMIs of embedding of TLDs from humanPFC cells in Lee’s ^9^data (560 cells).

\# means P < 0.05, * means P < 0.001, paired *t*-test for samples.

**Figure S4.** Assessment of deDoc2 with simulated data, and experimental scHi-C data.

a. Number of TLDs from simulated scHi-C data with threshold D=500 and simulated bulk Hi-C data.
b. Number of TLDs from simulated scHi-C data with threshold D=500 after imputation.
c. Structural entropy of predicted TLDs in Nagano’s data.
d. Modularity of predicted TLDs in Nagano’s data.
e. Fold change of ChIP-seq peak signals (CTCF, H3K4me3 and H3K36me3) between TLD boundaries and background (regions away from boundaries). TLDs are called from Nagano’s data. Error bars represent 95% confidence interval.

**Figure S5.** Assessment of deDoc2, deDoc, and Spectral TAD with downsampled data.

a. TLD_adjR2 standard error at 0-40kb of nested TLDs predicted by deDoc2, deDoc, and Spectral TAD in three downsample rates.
b. Number of nested TLDs predicted by deDoc2, deDoc, and Spectral TAD in three down-sample rates.
c. Length of nested TLDs predicted by deDoc2, deDoc, and Spectral TAD in three down-sample rates.
d. Length of nested sub-TLDs predicted by deDoc2, deDoc, and Spectral TAD in three downsample rates.

**Figure S6. APOE expression level in different celltypes.**

APOE expression level in Astro, MG,L23 and Vip cells.

**Figure S7. Assessment of deDoc2, IS and deDoc with bulk Hi-C data from Rao’s data.**

a. Number of TLDs from 6 cell type bulk Hi-C.
b. Length of TLDs from 6 cell type bulk Hi-C.
c. Fold change of ChIP-seq peak signals (CTCF, H3K4me3 and H3K36me3) between TLD boundaries and background (regions away from boundaries). TLDs are called from 6 cell type bulk Hi-C.
d. Modularity of predicted TLDs from 6 cell type bulk Hi-C.
e. Structural entropy of predicted TLDs from 6 cell type bulk Hi-C.
f. Contact density of predicted TLDs from GM12878 cell type bulk Hi-C.

**Figure S8. Comparison between deDoc2 and SuperTAD.**

a. Running time used for predicting TLDs of downsampled Hi-C from Rao’s data and experimental scHi-C from Tan’s data, 9 chromosomes total (chr14 to chr22).
b. Running time used for predicting TLDs from simulated data.
c. AMI between TLDs from simulated single-cell Hi-C at different simulation thresholds and bulk Hi-C.
d. WS between TLDs from simulated single-cell Hi-C at different simulation thresholds and bulk Hi-C.
e. Fold change of ChIP-seq peak signals (CTCF, H3K4me3 and H3K36me3) between TLD boundaries and background (regions away from boundaries). TLDs are called from downsampled Hi-C from Rao’s data and experimental single-cell Hi-C in Tan’s data in 9 chromosomes total (chr14 to chr22).

**Figure S9. Treatment of isolated vertices in Hi-C data.**

a. Example of unmappable regions potentially recognized as gap regions.
b. Example of isolated vertices potentially placed inside TLDs.
