## Supplementary figures and images for "DeDoc2 identifies and characterizes the hierarchy and dynamics of chromatin TAD-like domains in the single cells"

### Supplementary Figure 1

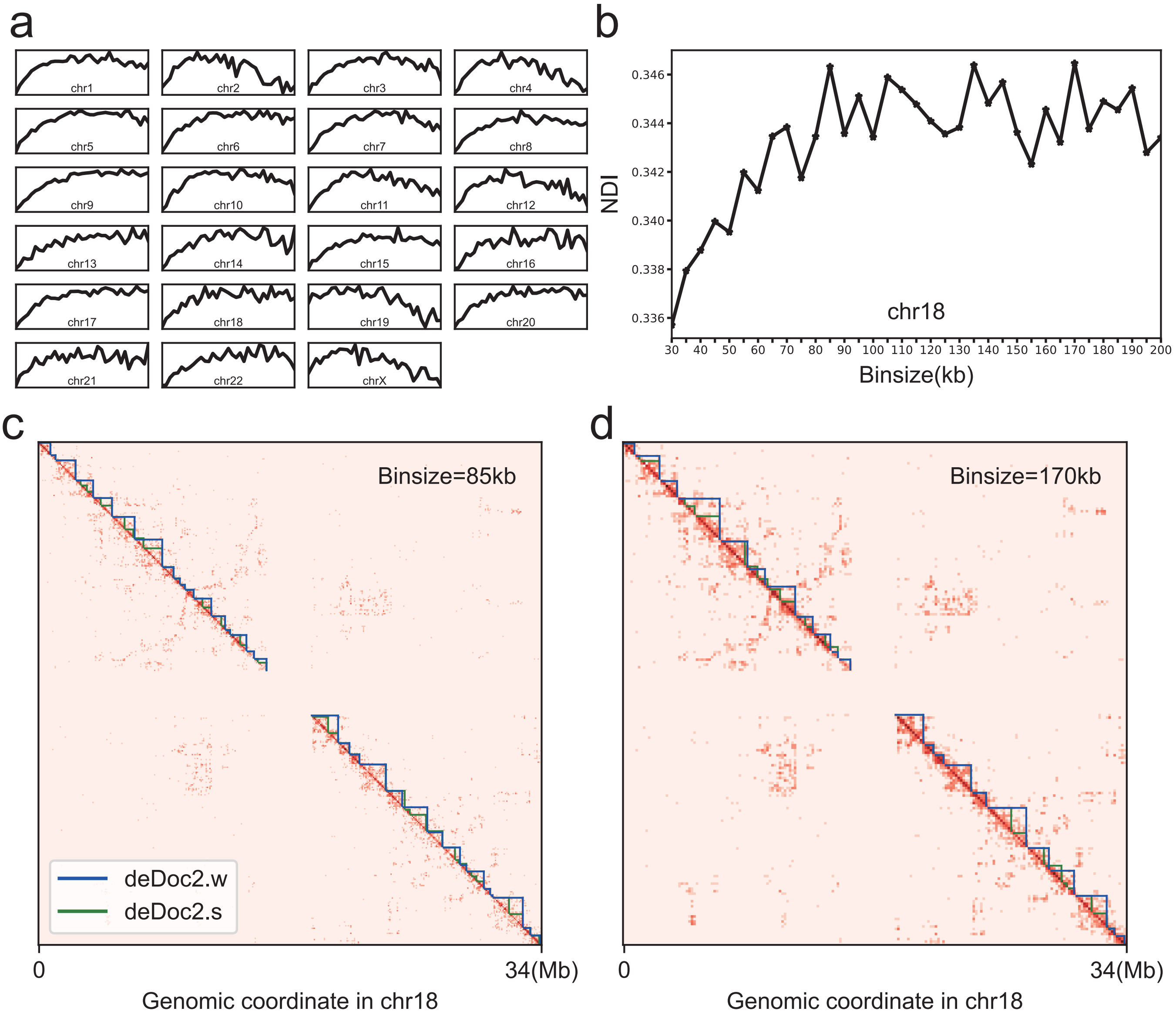

### Supplementary Figure 2

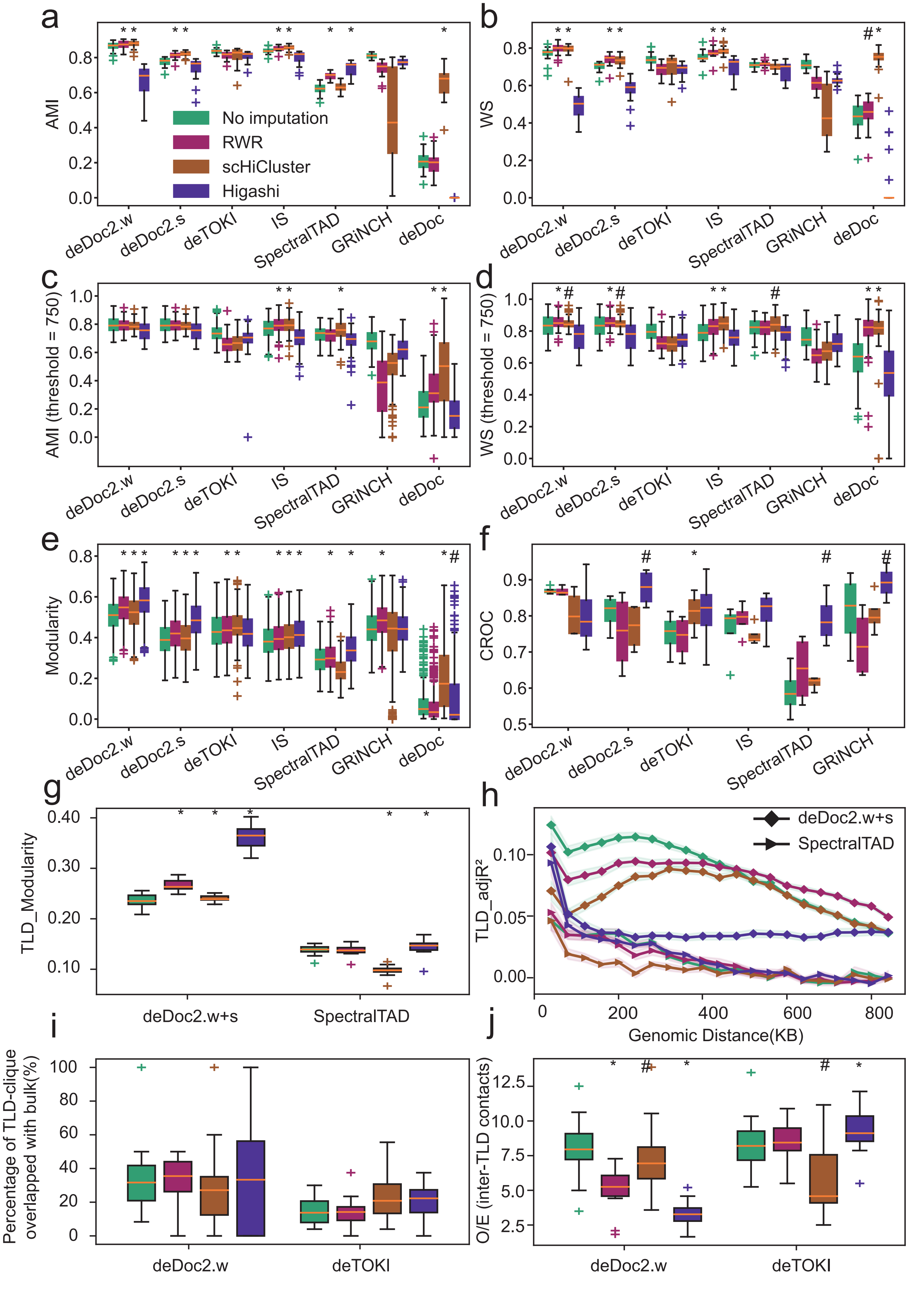

### Supplementary Figure 3

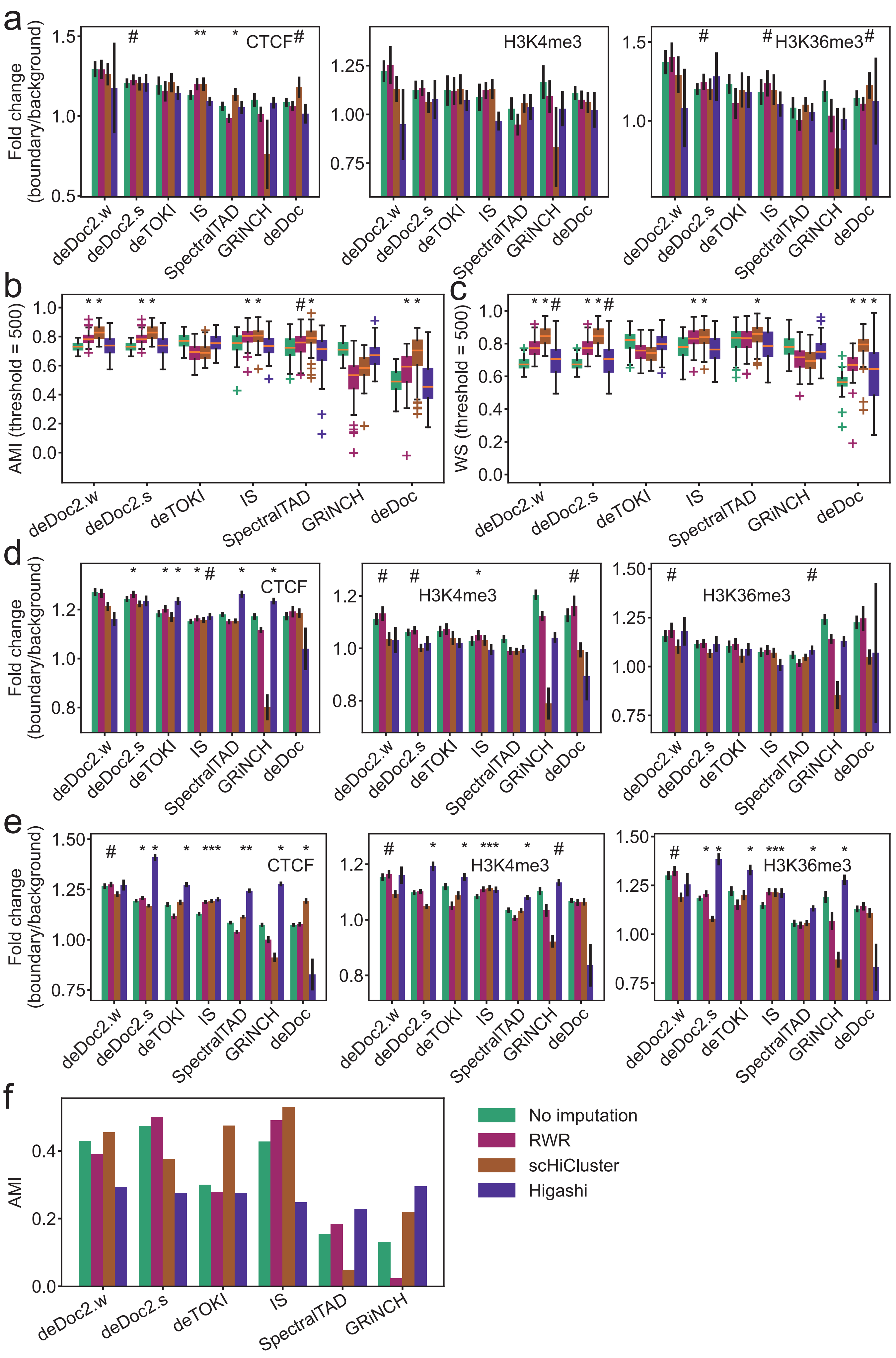

### Supplementary Figure 4

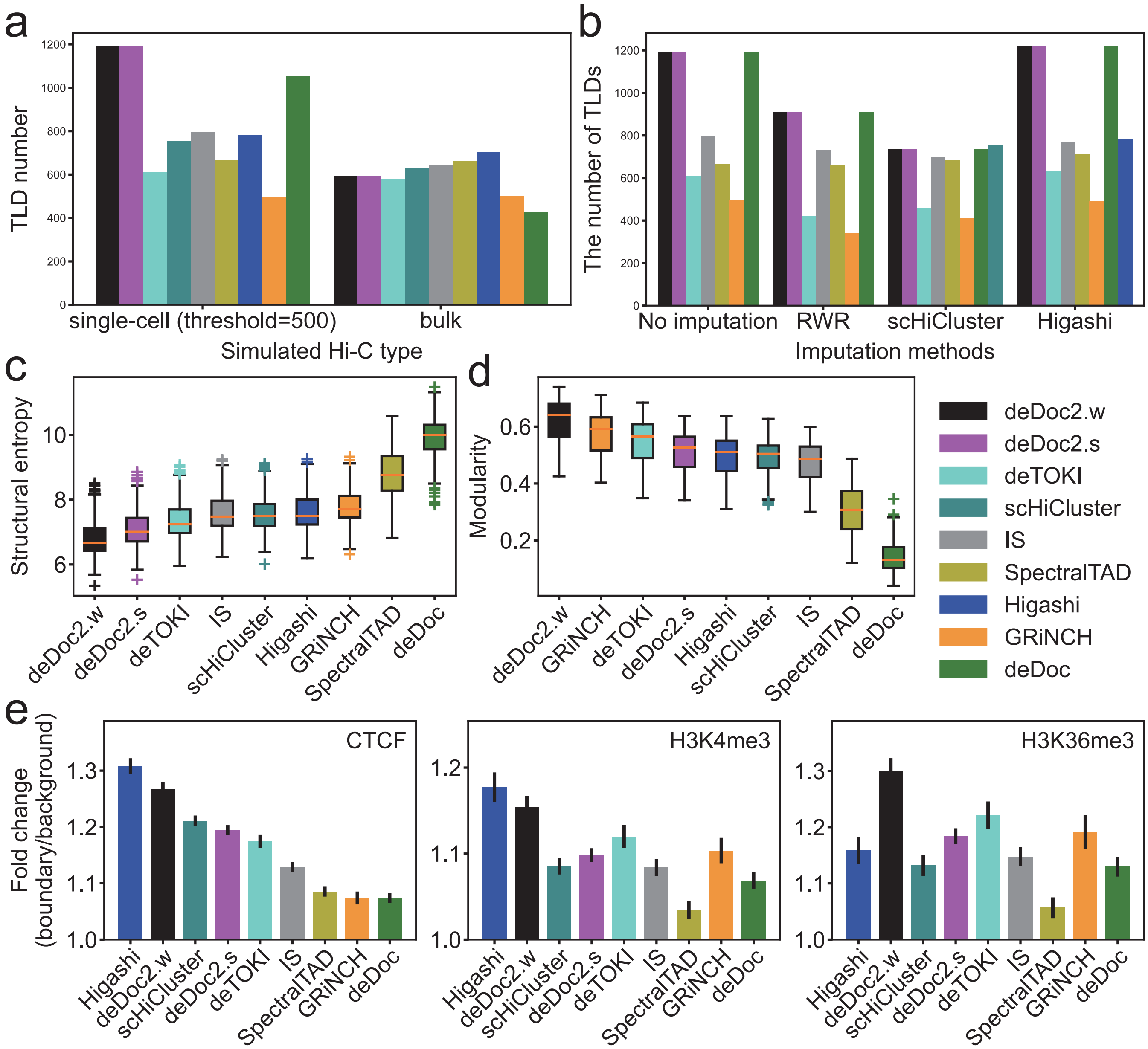

### Supplementary Figure 5

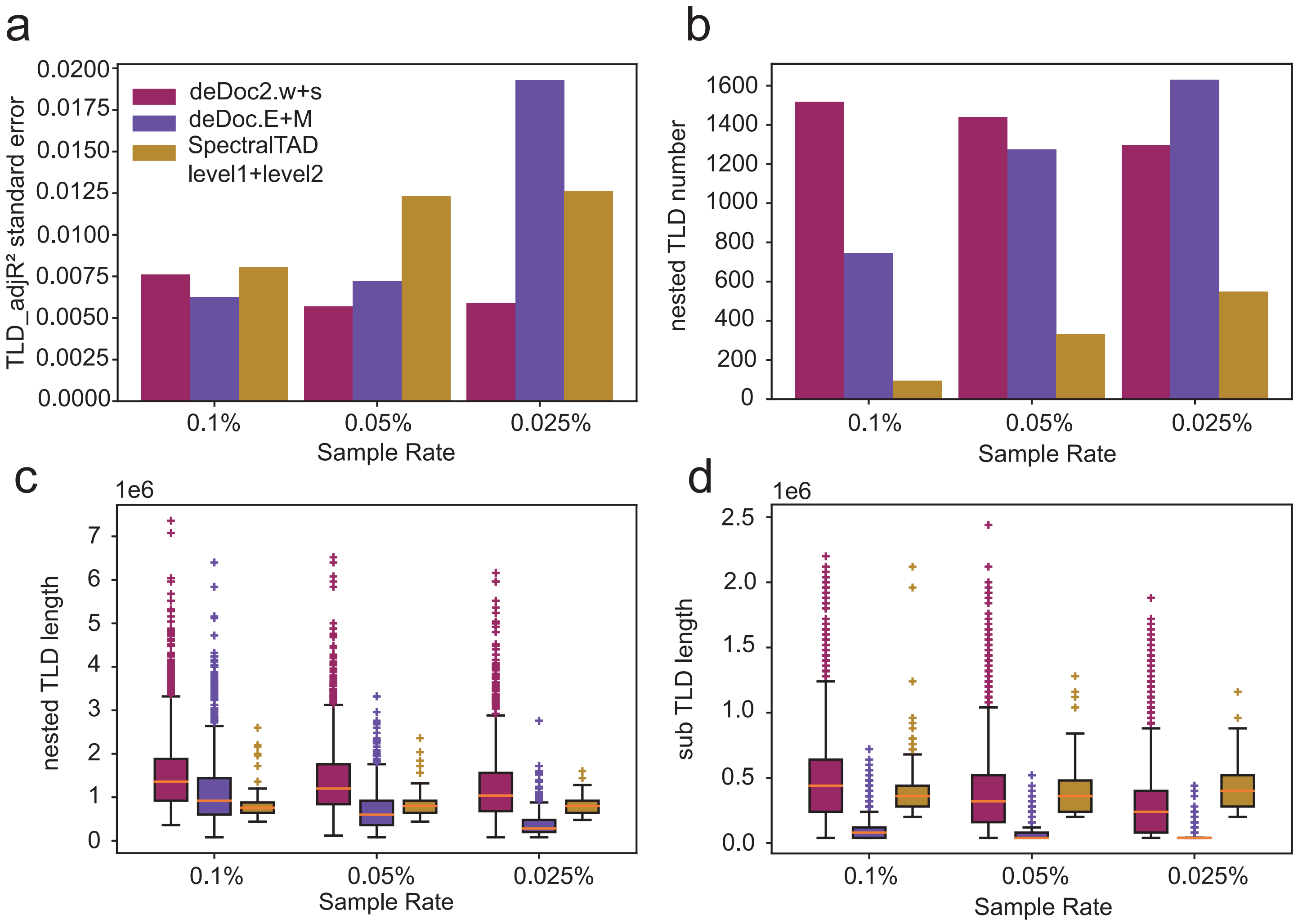

### Supplementary Figure 6

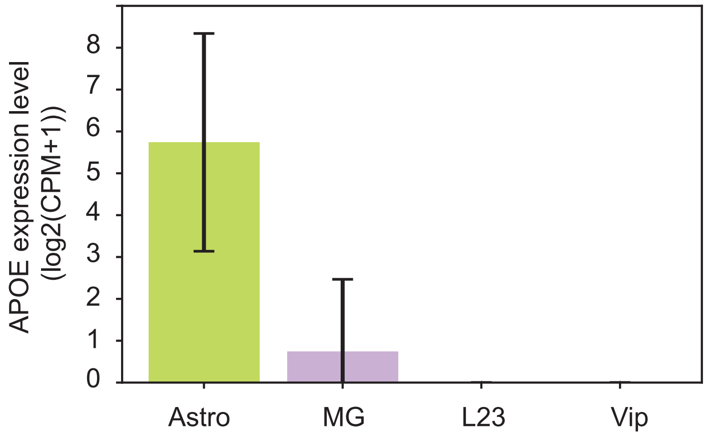

### Supplementary Figure 7

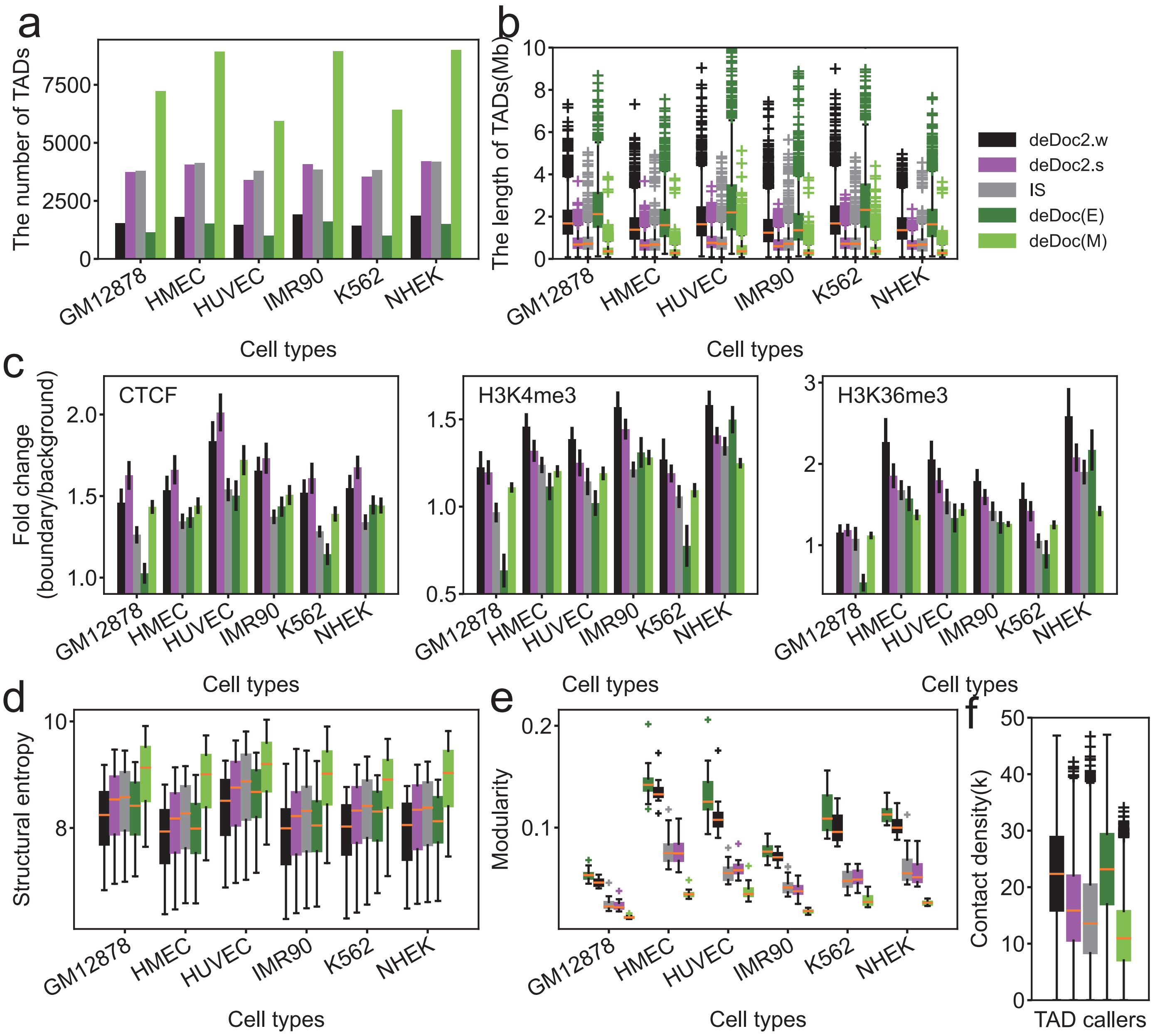

### Supplementary Figure 8

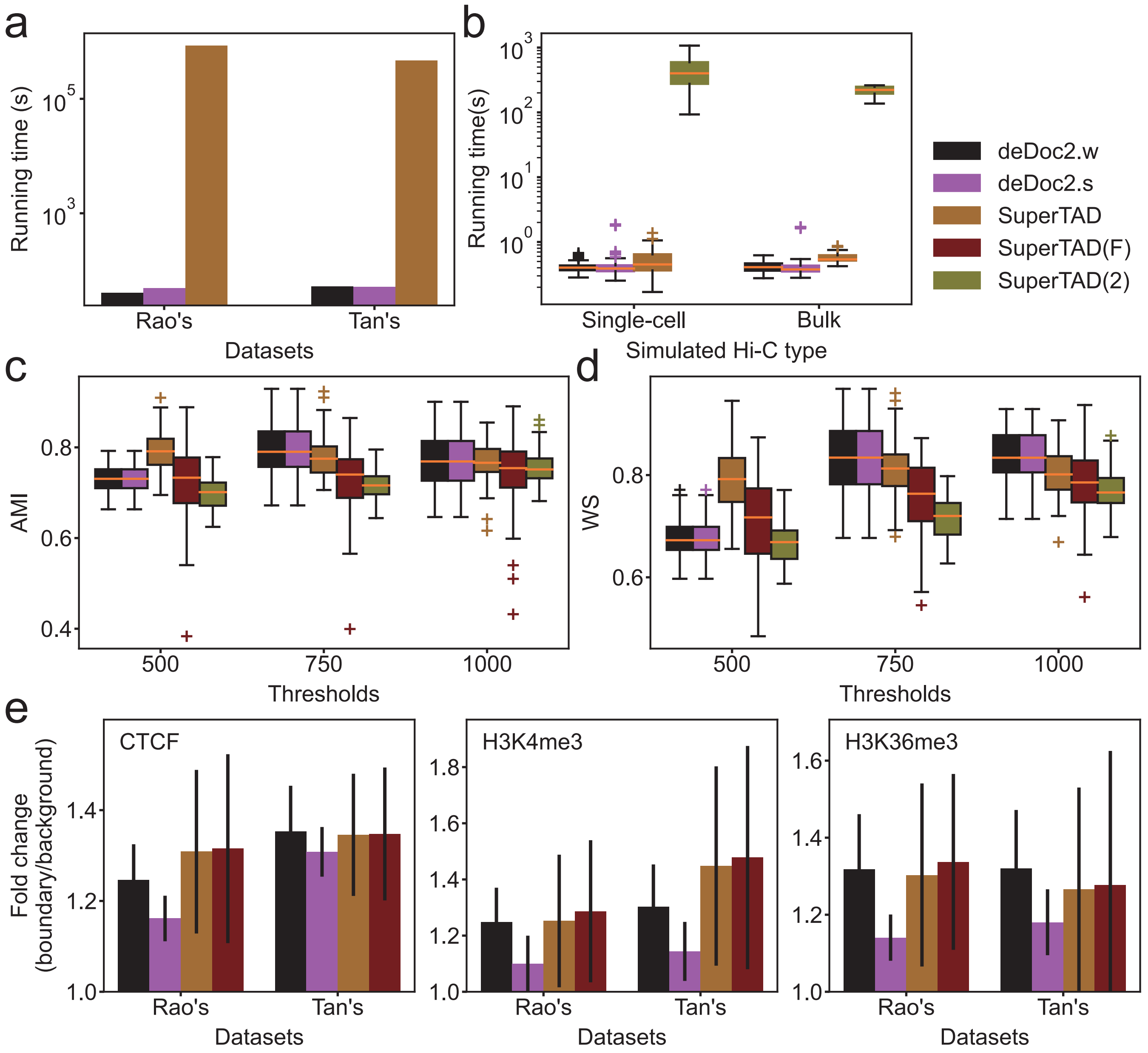

### Supplementary Figure 9

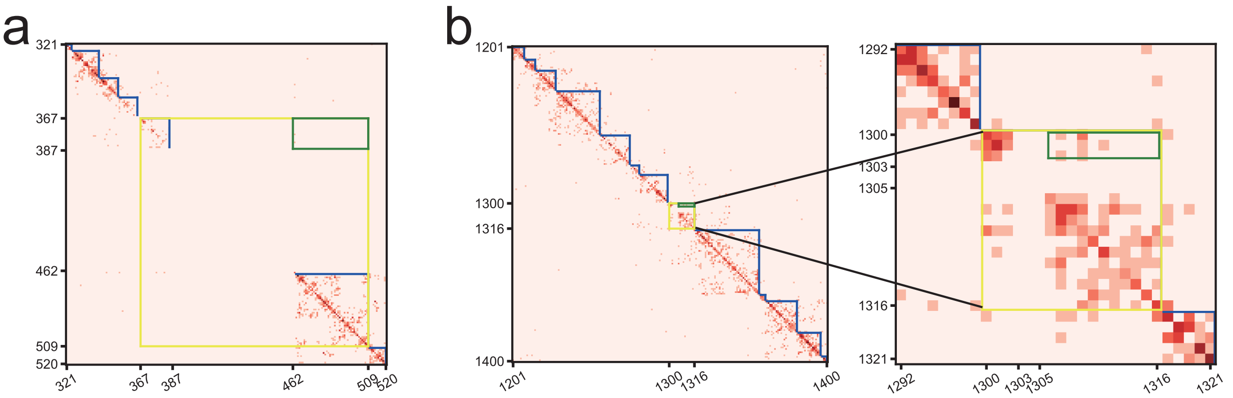
