## SupplementaryTable 4 for "DeDoc2 identifies and characterizes the hierarchy and dynamics of chromatin TAD-like domains in the single cells"

Table 4. Comparison of CPU running times between deDoc2 and SuperTAD.

| Tools | deDoc2.w | deDoc2.s | SuperTAD | Tools | deDoc2.w | deDoc2.s | SuperTAD |
| --- | --- | --- | --- | --- | --- | --- | --- |
| downsampled IMR90 cell from Rao’s data | | | | GM12878 cells from Tan’s data | | | |
| chr14 | 6.54s | 7.65s | 4771min | chr14 | 7.88s | 8.12s | 1594min |
| chr15 | 6.10s | 6.12s | 4266min | chr15 | 7.80s | 6.40s | 1445min |
| chr16 | 5.66s | 5.84s | 1572min | chr16 | 6.80s | 6.37s | 848min |
| chr17 | 4.29s | 5.74s | 1239min | chr17 | 5.68s | 5.92s | 1302min |
| chr18 | 4.24s | 5.52s | 881min | chr18 | 5.42s | 5.19s | 1076min |
| chr19 | 3.66s | 5.04s | 801min | chr19 | 4.96s | 5.15s | 645min |
| chr20 | 3.47s | 4.52s | 343min | chr20 | 4.87s | 4.87s | 381min |
| chr21 | 3.29s | 4.40s | 191min | chr21 | 4.50s | 4.79s | 194min |
| chr22 | 3.05s | 3.90s | 150min | chr22 | 4.42s | 4.56s | 335min |

Note: All tools were tested using 1 CPU core. The results of SuperTAD(F) can be calculated the same time as SuperTAD. Chromosomes were not segmented before put into SuperTAD (further discussion can be found in supplementary text).

CPU: Intel(R) Xeon(R) Gold 6240 CPU @ 2.60GHz
