## SupplementaryTable 5 for "DeDoc2 identifies and characterizes the hierarchy and dynamics of chromatin TAD-like domains in the single cells"

Table 5. CPU/GPU running times with single-cell Hi-C of GM12878 cells from Tan’s data.

| Tools | Time (min) | Remark |
| --- | --- | --- |
| deDoc2.W | 55.57 |  |
| deDoc2.S | 64.64 |  |
| deTOKI | 322.16 |  |
| scHiCluster | 529.02 | scHiCluster imputes scHi-C data and use TopDom to predict TLDs from imputed Hi-C data. We calculated both the time for imputation and TLDs prediction by TopDom. |
| IS | 1242.58 |  |
| SpectralTAD | 301.04 | We calculated the time with parameter “levels=1” which did not predict hierarchical structures. |
| Higashi | 110.42  (1 GPU & 1 CPU core) | Higashi imputes scHi-C data and uses insulation score based method to predict TLDs from imputed Hi-C data. We calculated both the time for imputation and TLD prediction. We set the parameter nbr=0 as author suggested. |
| GRiNCH | 132.42 |  |
| deDoc | 30.59 |  |

Note: All tools were tested using 1 CPU core, except for Higashi which requires GPU and we tested using 1 GPU with 1 CPU core.

CPU: Intel(R) Xeon(R) Gold 6240 CPU @ 2.60GHz

GPU: Tesla V100 SXM2 32GB
